# Cellular deconvolution of the brain with topological magnetic resonance image analysis

**DOI:** 10.64898/2025.12.10.693426

**Authors:** Luis A. Vazquez, Michael B. Fromandi, Tracy L. Hagemann, Ryan D. Risgaard, Jose M. Guerrero-Gonzalez, Ajay P. Singh, Paloma C. Frautschi, Samuel A. Hurley, André M. M. Sousa, Doug C. Dean, Tyler K. Ulland, John-Paul J. Yu

## Abstract

Magnetic resonance imaging (MRI) is foundational tool in neuroscience, enabling characterization of neuroanatomical markers of disease, behavior, and cognition. However, the precise cellular processes driving the structural and functional readouts provided by MRI remain opaque. Non-invasively assessing cell type, abundance, and location using MRI has the potential to revolutionize both basic science and clinical practice. To this end, we developed <u>S</u>pa<u>T</u>ial <u>R</u>epresentation and <u>A</u>nalysis using <u>T</u>opological <u>A</u>rchitecture (STRATA), an image-based gradient-boosted machine learning framework, which quantifies cell type proportions of neurons, astrocytes, oligodendrocytes, and microglia from MR images. Here we demonstrate and validate STRATA on diverse disease models, species, and regions of interest that together highlight the generalizability of the STRATA framework.

## Main

Magnetic resonance imaging (MRI) is one of the most powerful noninvasive tools for studying the living brain, enabling detailed characterization of neural architecture, functional networks, and white matter pathways with high spatial and temporal resolution. Despite its remarkable contributions, MRI remains fundamentally a macroscopic technique. Due to inherent technical and spatiotemporal constraints, MRI can only indirectly measure neuropathology by generating imaging phenotypes that serve as nonspecific proxies for the molecular and cellular processes underlying pathological changes in the brain (*1, 2*). Because MRI-derived imaging phenotypes ultimately reflect the underlying composition of neurons and glial cells such as astrocytes, microglia, and oligodendrocytes, the absence of imaging or analytical methods capable of directly characterizing these cellular populations or resolving their relative abundance constrains the ability of MRI to uncover causal neurobiological mechanisms of disease. As a result, clinicopathologic insights from modern neuroimaging remain largely correlational and without a deeper understanding for how cellular and molecular processes are encoded in MR images, our capacity to derive mechanistic insight into brain function and pathology *in vivo* with MRI will continue to be limited.

This limitation has motivated the development of quantitative MRI (qMRI) techniques designed to move beyond qualitative image contrast and toward metrics that more directly quantify the molecular and cellular composition of tissue. By quantifying intrinsic tissue properties like intrinsic relaxation (T_1_-, T_2_-mapping), perfusion and flow (arterial spin labeling, dynamic contrast enhancement), or magnetic susceptibility/chemical composition (proton density fat fraction), qMRI aims to transform MRI from a descriptive imaging tool into a system capable of linking image features to specific biological mechanisms. In the context of neuroimaging, quantitative imaging with diffusion weighted MRI (dMRI) has shown promise for linking neurobiology to MRI as dMRI methods are uniquely sensitive to the microscopic organization and composition of neural tissue. Water diffusion in the brain is constrained by cellular structures such as membranes, axons, dendrites, and organelles and the ordered pattern and arrangement of these cellular components allows dMRI to capture information about cell density, size, and orientation. These features also directly correlate to the cellular composition of brain tissue itself and can provide a biologically interpretable bridge between dMRI measures and underlying cytoarchitecture.

Of the numerous dMRI techniques developed, neurite orientation dispersion and density imaging (NODDI) (*3*) has proven remarkably effective for its ability to bridge MRI imaging phenotypes and neurobiology. NODDI biophysically models water diffusion in brain tissue into intracellular or extracellular compartments that further refines the sensitivity of dMRI to unique microstructural environments in the brain. This enhanced sensitivity to brain cytoarchitecture is reflected in recent work demonstrating NODDI and similar multi-compartment diffusion models to be sensitive and directly related to the underlying cellular organization of the brain (*4–6*) and for its exceptional sensitivity to water diffusion on the length-scale of neurons and glia (*7–9*). Moreover, our recent work demonstrated that spatial patterns of gene expression in the brain corelate with intracellular and extracellular water diffusion in the NODDI model (*2*), making NODDI one of the most sensitive and direct imaging representations of neurobiology. Building on the relationship between NODDI dMRI and cellular architecture, we sought to develop an approach for quantifying cellular composition with MRI. We drew inspiration from a central hypothesis in neuroscience that the spatial organization of brain tissue, or cytoarchitecture, is defined by the number, type, and density of neurons and glial cells present (*10, 11*). Correspondingly, the spatial organization of signal intensity present in a MR image must also be fundamentally correlated to the number and type of cells present in the image (e.g., MR images of white matter regions correlate with higher oligodendrocyte cell counts than in gray matter) (*12*). Provided these correlations, we hypothesized that spatial quantification of signal intensity in a MR image, which is a direct representation of underlying brain cytoarchitecture, can therefore provide inferences into the number and types of cells present in the image (**Fig. 1A**).

**Fig. 1.**
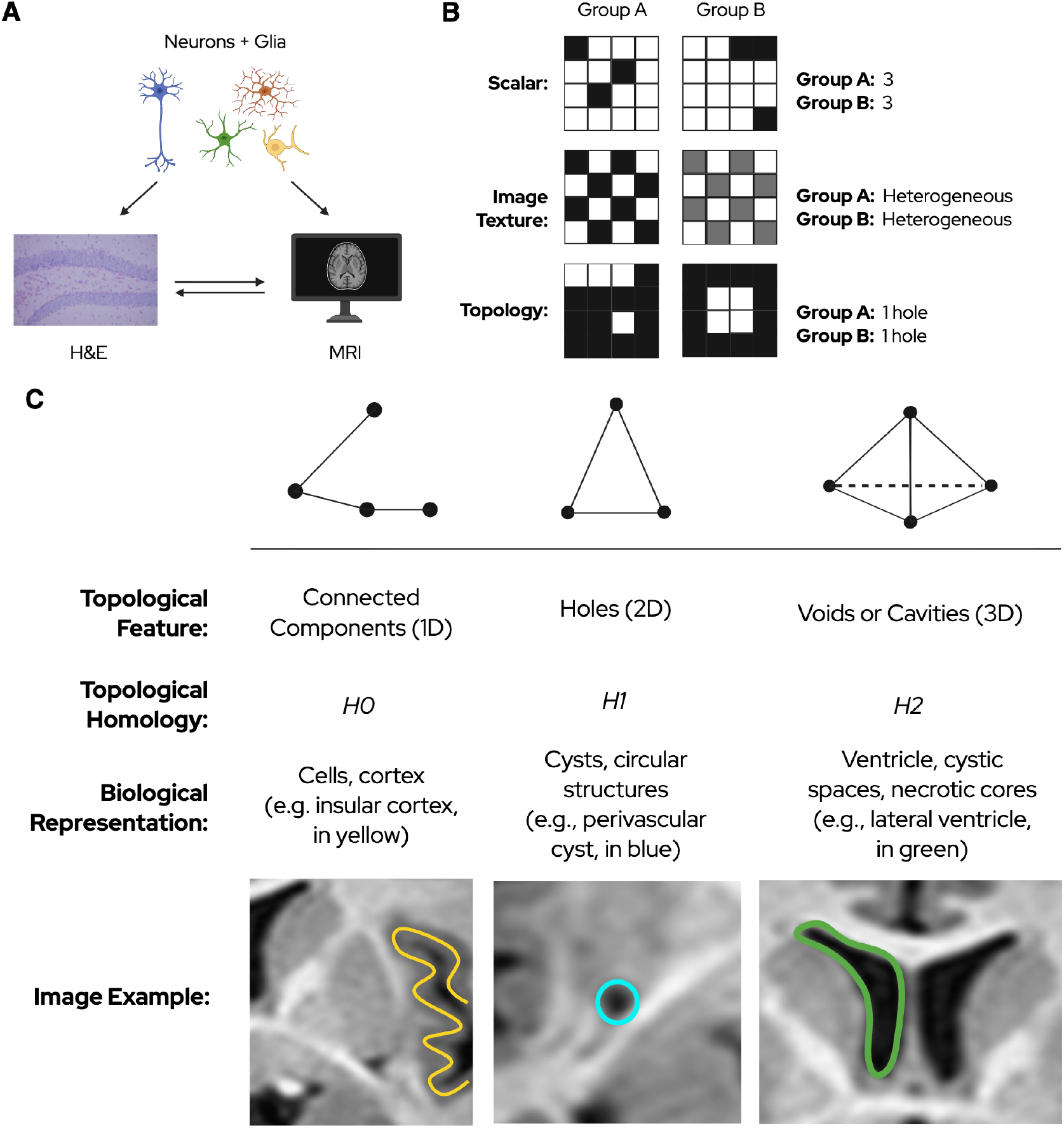
**(A) Hypothesis and rationale.** Individual neurons and glial cells (top) spatially organize into characteristic cytoarchitectural patterns that are defined by the number, type, and density of neurons and glial cells present (histology, bottom left). Similarly, the spatial organization of signal intensity present in a MR image is also fundamentally correlated to the number and type of cells present in the image (MRI, bottom right). Provided these correlations, we reasoned that spatial quantification of signal intensity in a MR image, which is a direct representation of underlying brain cytoarchitecture, can therefore provide inferences into the number and types of cells present in the image. **(B) Image analysis methods in qMRI**. Scalar approaches to image analysis are insensitive to the spatial distribution of voxel intensities in a predetermined region of interest. Here, the scalar value of Group A and B is 3. Each group, however, contains a different spatial pattern of signal intensity and thus would be non-specific to processes giving rise to the different signal distribution despite the having the same overall scalar value. Similar non-specificities are also seen in image texture analysis. As an illustration, Groups A and B possess the identical pixel distributions but with different overall intensity that would give rise to different calculated gray level texture features. However, as gray level texture features can be affected by image acquisition or reconstruction parameter differences (e.g., coils, field strength), understanding whether these differences result from differences in technique or underlying pathology reinforce both the non-specificity of image texture approaches as well as their low reproducibility and interpretability. In contradistinction, topological approaches instead quantify the ‘shape’ of the data by quantifying image features such as ‘holes’ as shown. These features are topologically invariant (e.g., these features remain the same even following image deformation, such as stretching, bending, or twisting), which contributes to the robustness and reproducibility of topological analyses. Here, despite significant differences in pixel number and distribution, Groups A and B both topologically possess just 1-hole and are thus quantified as being equal. **(C) Topological features present in MR images**. (Left column) The topological group H0 represents connected components, which are 1-dimensional structures such as points and edges that do not enclose any space. In brain MR images, H0 components correspond to distinct cellular regions with uniform signal intensity, such as the insular cortex (highlighted in yellow), where a continuous cortical mantle creates a connected component separate from surrounding structures. (Middle column) The topological group H1 represents holes, which are 2-dimensional spaces that are enclosed by a single connected component. These topological features capture circular or ring-like structures in the image plane. In brain MR images, H1 components correspond to structures such as perivascular cysts (highlighted in blue), where fluid-filled spaces create circular voids surrounded by tissue. (Right column) The topological group H2 represents voids or cavities, which are 3-dimensional spaces that are completely enclosed by a connected component. These features capture volumetric cavities within the brain. In brain MR images, H2 components correspond to large structures such as the ventricles (highlighted in green), where cerebrospinal fluid creates a 3D void surrounded by brain parenchyma. Each topological group captures progressively higher-dimensional topological features, enabling comprehensive characterization of brain tissue organization.

To quantify spatial patterns in imaging data, recent attention has turned to topological data analysis (TDA), which leverages tools from algebraic topology to measure geometric structures present at both local and global scales (*13*). Quantitative image analysis conventionally performs analysis at the pixel level with summaries of mean pixel intensity in a region of interest (ROI) or through the generation of texture-based statistics that describe local signal variation or local image features (**Fig. 1B**). Because pixel level analyses treat voxels as independent observations, they fail to capture larger spatial patterns of signal intensity that are present in brain tissue, how they are spatially organized, and how different regions relate or are delineated from each other. In contrast, image analysis with TDA instead focuses on the spatial organization of signal intensity in the image (or imaging volume) rather than the absolute intensity values themselves or local voxel statistics. TDA considers an imaging ROI as a single geometric volume and enumerates the multidimensional shapes formed by the data. This is done by connecting neighboring voxels based on both their spatial proximity and intensity similarity (see *Supplementary Material* for a full mathematical formulation and discussion). This provides information on patterns such as connected components (1D), holes (2D), and cavities (3D) that are present in the data, which reflect the multidimensional cytoarchitecture of brain tissue (**Fig. 1C**). Furthermore, because the emphasis of TDA is on *relative* patterns of signal intensity instead of *absolute* values of signal intensity, TDA analyses sidestep several unresolved challenges in qMRI including hardware, scanner, and sequence variability, image noise, and fitting instability, making TDA inherently more generalizable and reproducible across sites, scanners, and acquisition protocols (*14, 15*).

To harness the biological sensitivity of NODDI dMRI and the analytic advantages of TDA, we have developed STRATA (<u>S</u>pa<u>T</u>ial <u>R</u>epresentation and <u>A</u>nalysis using <u>T</u>opological <u>A</u>rchitecture), a machine learning–based image analysis pipeline designed to infer relative group differences in cell type proportions of neurons, astrocytes, oligodendrocytes, and microglia directly from MRI data. STRATA is trained using topological embeddings extracted from NODDI imaging data with matched histologically quantified regions to learn nonlinear mappings between topological features and cellular abundance using gradient boosting (**Fig. 2**). Once trained, STRATA applies the learned model to unseen MRI data to infer relative group differences cell-type proportions, enabling quantitative estimation of tissue composition in the absence of histological data. Here, we demonstrate the sensitivity, specificity, and reproducibility of STRATA to estimate cell-type proportions across a diverse range of disease models spanning neurodevelopmental to neurodegenerative disorders from MRI data alone. We further showcase the ability of STRATA to deconvolve cellular populations in multiple brain regions in murine, rat, and non-human primate models that further demonstrate the robustness and trans-species generalizability of the STRATA model.

**Fig. 2.**
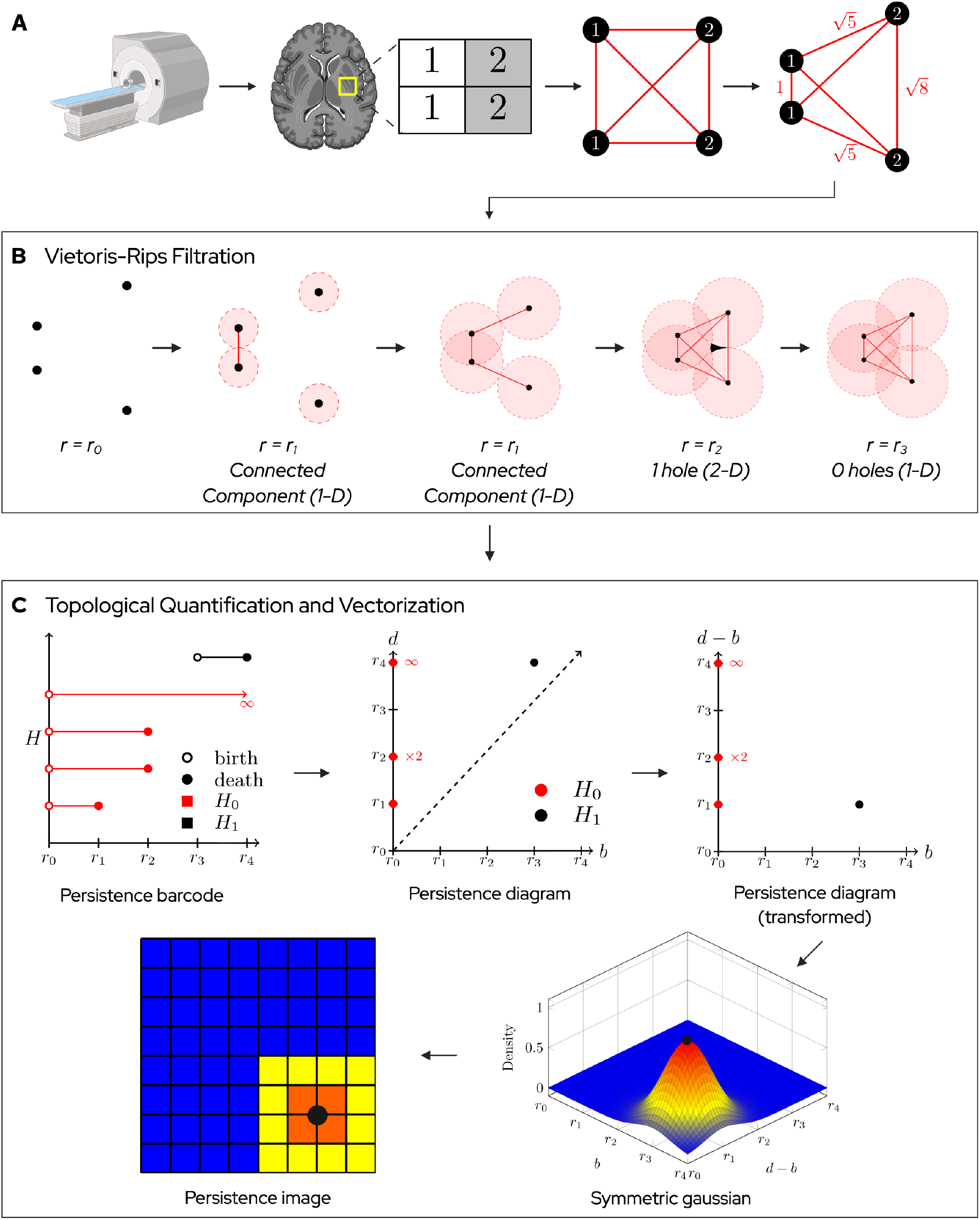
Topological data analysis of MR images. **(A) Point cloud embedding of imaging data.** Following qMRI acquisition, the image volume is parsed into its constituent voxels, with each voxel assigned its associated quantitative metric. For the purposes of illustration, we will consider a reduced 2x2 pixel grid of a 2D slice. Here, each voxel is first embedded as a spatial graph, where individual pixels are converted into nodes and adjacent nodes are connected through edges where the length of each edge is defined by the Euclidean distance between nodes, which by default, is 1 for neighboring nodes. A weighted Euclidean distance between nodes is next calculated by incorporating the intensity of each pixel as a weighted in the calculated distance (see *Materials and Methods for a full mathematical formulation*). This weighted Euclidean distance between nodes causes edges between adjacent nodes with dissimilar intensities to increase with the result having both spatial and intensity-based differences between all pixels embedded into a mathematically equivalent representation. (**B**) **Illustration of the Vietoris-Rips Filtration**. To formally calculate the topology of these nodes and their spatial relationships now defined by their weighted Euclidean distances, we use the Vietoris-Rips filtration. This process involves building circular neighborhoods around each node and progressively increasing the radii, drawing edges between nodes once neighborhoods contact another. At r_0_, we have neighborhoods of radii 0, and thus no nodes are connected. As we increase the radius to some radius [r_1_], the neighborhoods of the nodes on the left and right side of the rectangle contact each other and the two edges are drawn. As we continue to increase the radius, at some larger radius [r_2_], the neighborhoods of the node edges on the top and bottom of the rectangle contact and the top and bottom edges are drawn. This now creates a loop between all the nodes, generating a hole in the middle where the neighborhoods do not overlap (filled in as black). By increasing the radii further to some radius [r_3_], the adjacent neighborhoods overlap and cover the hole. (**C**) **Topological feature quantification and vectorization**. This process by which holes appear and disappear (and for 3D imaging volumes, voids and cavities appear and disappear) is tracked as a function of the size of the radius of the nodes, which can be graphically displayed as a persistence barcode where the appearance (and disappearance) of features like holes are tracked as births and deaths. On the barcode, H0 represents the birth of all nodes (also called connected components) and thus are represented at r_0_. In our example, there are 4 H0 components since we have 4 nodes corresponding to each pixel. As shown in panel **B** above, at some radius r_1_, we drew edges on the left and right side of the rectangle with these individual nodes now merging into a single component, we thus went from having two connected components (two nodes) to just one, thereby causing one connected component (e.g., node) to die. A similar merging occurs on the right side of the rectangle leaving just two connected components alive overall. At our larger radius r_2_, we drew the top and bottom edges, merging the two components representing the right and left edges into one. The surviving connect component represent a loop which also generates a 2D hole, giving birth to an H1 component (hole). At our larger radius r_3_, this hole is filled as it becomes completely covered by the surrounding radii, killing the H1 component (the hole). However, a single component persists at r_2_ and r_3_, first representing the square loop and then the covered square. By mapping birth time to an x-axis, and death time to a y-axis, we can create a persistence diagram representing the same information. Notice that since all H0 components are born at 0, they all appear on the y-axis. Additionally, as birth must occur prior to death, no topological components can appear below the x=y diagonal. For convenience, we next linearly transform the y-axis of the diagram to represent death-birth, such that the same information now lies on a square coordinate plane (persistence diagram, transformed). To vectorize this information for statistical analysis, we transform the transformed persistence diagram into a probabilistic representation. This is done by generating a symmetrical gaussian around each component, representing the density of points in that region of the transformed persistence diagram. In this figure, we illustrate the symmetric gaussian around the H1 component. We can then integrate across this surface in equidistant bins to create a new matrix representation called a persistence image. Each bin is then treated as an individual feature for downstream machine learning.

### Generation and validation of the STRATA model

The STRATA model is based on the premise that spatial patterns of MR signal intensity can provide insights into the number and types of cells present. The rationale behind STRATA can be understood by considering the MR signal pattern of white and gray matter structures, such as the corpus callosum and thalamus. In each case, the distinct MR signal pattern (e.g., T1-weighting) of these areas possesses strong correlation to their corresponding cellular composition (primarily oligodendrocytes in the corpus callosum and neurons in the thalamus). With STRATA, we explored whether this rationale could be extended to other cell types and other areas of the brain with greater cytoarchitectural complexity. To train the STRATA model, we used a newly developed rat model of Alexander disease (AxD) (*16*) (**Fig. 3A**) The AxD model demonstrates a broad dynamic range of neuronal and glial pathology (*17*), documented sex-specific changes, and substantial quantitative changes present on NODDI imaging (*16, 18*) that together comprise wide-ranging imaging and cellular features to impute into a machine learning model. The STRATA training dataset includes two cohorts of male and female outbred control and AxD animals. NODDI imaging was performed in one cohort (Cohort 1), and bulk tissue transcriptomics of the right hippocampus were performed in a second independent cohort (Cohort 2). NODDI images from Cohort 1 were normalized and registered to a common template. The right hippocampus was then masked and extracted, and topological features calculated from neurite density index (NDI) and orientation dispersion index (ODI) NODDI parameter maps. For Cohort 2, cell-type composition from previously published bulk tissue RNA-seq data of the right hippocampus of AxD animals (*19*) was estimated using dampened weighted least squares (DWLS) (*20*) (**fig. S1**).

**Fig. 3.**
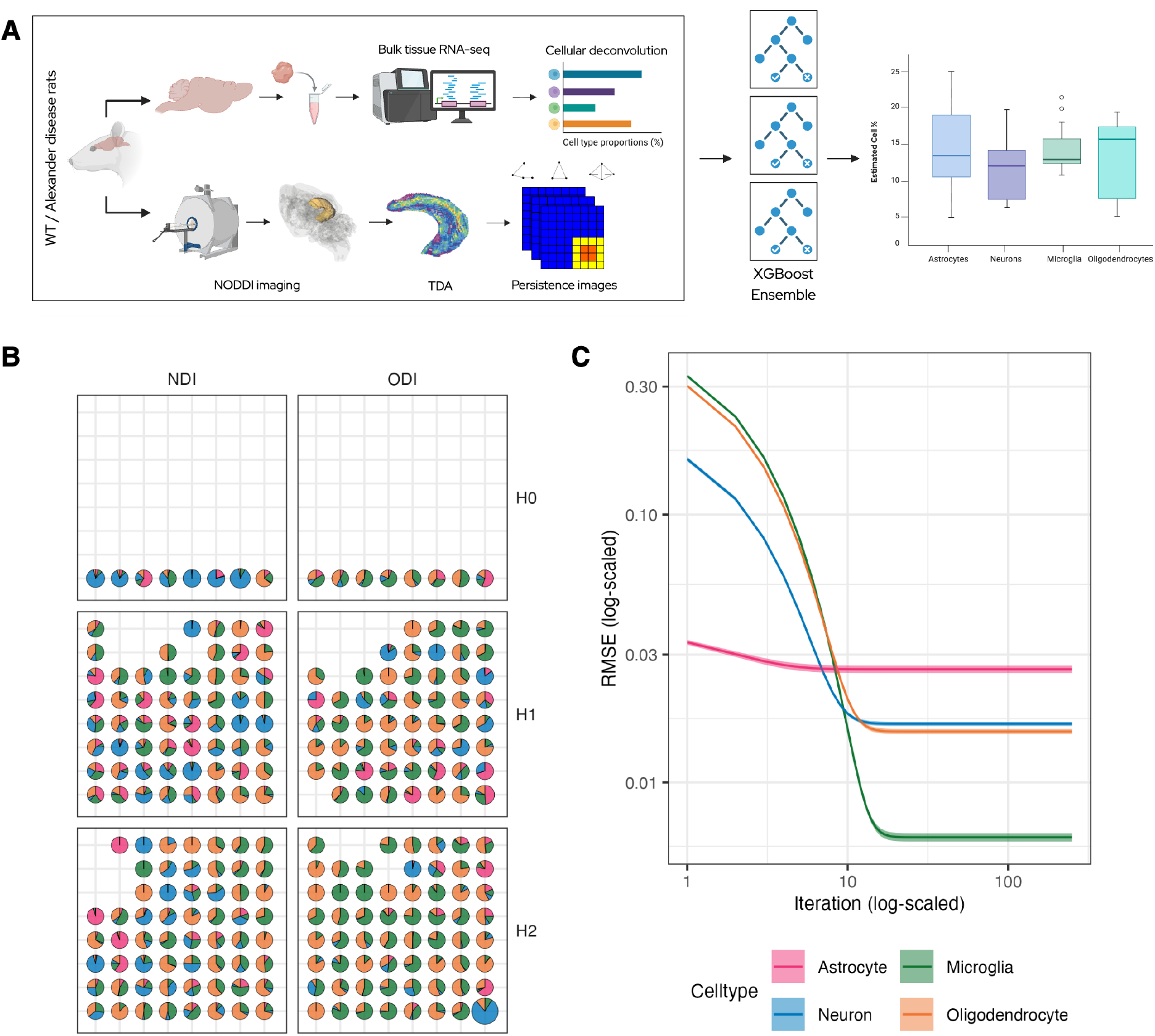
Overview of the STRATA machine learning pipeline. (**A**) To train the STRATA model, two cohorts of male and female outbred control rats and Alexander disease animals were used. NODDI imaging was performed in one cohort (Cohort 1), and bulk tissue transcriptomics of the right hippocampus were performed in a second independent cohort (Cohort 2). For the imaging cohort, multicompartment diffusion weighted imaging (MC-DWI, NODDI) was used to generate neurite density index (NDI) and orientation dispersion index (ODI) maps, which are embedded as weighted spatial graphs. A representative right hippocampus map is shown, as this was the ROI used for training. TDA was applied to extract features in the form of persistence images, producing three arrays per imaging modality corresponding to homological dimensions H0, H1, and H2. For transcriptomics, the right hippocampus, including Ammon’s horn and the dentate gyrus were used for bulk tissue RNA-seq, which was subsequently deconvolved using a reference single cell RNA-seq dataset via the Dampened Weighted Least Squares (DWLS) method, yielding estimated cell type proportions for each sample. After deconvolution, subtypes of major cell types (astrocyte, microglia, neuron, and oligodendrocytes) were grouped together. Persistence image features were then paired with the deconvolved cell type proportions, which are treated as ground truth labels. An ensemble of XGBoost models was then trained to estimate individual cell type percentages from imaging features. (**B**) **Mean SHAP values for persistence image features**. To evaluate the performance of the trained STRATA model, we constructed a grid of plots showing the mean SHAP (SHapley Additive exPlanation) values for features derived from persistence image. Each plot corresponds to the persistence image of a specific modality (columns) and homological dimension (rows). Within each plot, the x-axis represents birth time and the y-axis persistence, corresponding to bin positions in the 8×8 persistence image. Each pie chart represents a single persistence image feature and shows its mean contribution to the prediction of different cell types, calculated across the ensemble of XGBoost models. Pie slice colors match cell types as indicated in the legend. The size of each pie chart reflects the overall importance of that feature, with larger charts indicating higher mean SHAP values. Bins without a pie chart represent features that were not used in the final models. Of note, these results demonstrate strong feature independence (low feature covariance) and robustness (stable predictions following bootstrapping) with excellent utilization of the topological feature space. (**C**) **Training RMSE**. The x-axis shows the number of boosting rounds (trees) during XGBoost training, and the y-axis shows the low root mean square error (RMSE) between successive iterations. Both axes are log-scaled for visualization. Each line represents the mean RMSE trajectory, computed across the ensemble of LOO models for a specific cell type. The line width denotes the standard deviation.

For model training, a bootstrapped permutation approach was used to pair data from Cohort 1 (imaging) and Cohort 2 (cell-type composition). Each rat from Cohort 1 was paired with all rats in Cohort 2 of both the same sex and the same genotype to create multiple paired samples while considering only biologically plausible pairings (**fig. S2**). Extracted topological features and cell-type composition data were then used to train separate models for each cell type (astrocytes, neurons, microglia, and oligodendrocytes) to test the efficacy of predicting each cell type in isolation (*21–23*) (**fig. S3**). A leave-one-out (LOO) approach was then implemented such that a separate model was created for each left-out subject from either cohort in which all permutation pairs, including that subject, were removed to reduce bias. Thus, many independent models were made for each cell type, and we relied on ensemble predictions from all LOO models to make groupwise comparisons using the Wilcoxon Rank Sum test (Bonferroni corrected). Model training was evaluated by examining variable importance features, correlation matrices, SHapley Additive exPlanation (SHAP) values (*24*), which together demonstrate strong feature independence (low feature covariance) and robustness (stable predictions following bootstrapping) with low training root squared mean error (RSME) indicating robust prediction accuracy and model fit (**Fig. 3B-C**).

### Testing of the STRATA model

As a proof-of-concept and positive control, we first sought to predict relative differences in cell type proportions for neurons, astrocytes, oligodendrocytes, and microglia in the left hippocampus between outbred wild type and AxD rats as model training used only data from the right hippocampus. Mirroring both results from the right hippocampus and prior published work (*16, 17*), STRATA correctly predicted the statistically significant relative differences in cell type proportions in all cell types with appropriate directionality (e.g., AxD rats have relative increased percentage of astroglia, microglia, and oligodendrocytes and decreased number of neurons in the hippocampus) (**Fig. 4A**). Importantly, STRATA correctly predicted the relative increase in number of oligodendrocytes present in the hippocampus of AxD rats compared to outbred wild type animals, which is consistent with prior published work, which showed transcripts for genes related to myelination such as *Plp1, Mag, Myrf*, and *Olig2* are slightly elevated in AxD rat hippocampus and show marginal enrichment in gene ontology analysis (*17*). To test the performance of the cellular sensitivity of the STRATA model, we next used STRATA to deconvolve relative cellular populations of microglia in a *Disc1* rat genetic model of psychiatric illness (*25*). In the *Disc1* model as in the brain, microglia are one the least prevalent major cell types present and thus an excellent barometer of the sensitivity of STRATA to measure relative changes in this smaller pool of cells. We have previously reported the effect of the antioxidant *N*-acetylcysteine (NAC) on microglia counts in the hippocampus of *Disc1* rats showing the relative increase in number of microglia present in *Disc1*-treated animals as validated by quantitative immunofluorescence (*26*). Using NODDI imaging data from these *Disc1* animals, STRATA correctly predicted the relative increased proportion of microglia in *Disc1*-NAC treated animals when compared to vehicle treated animals (**Fig. 4B**). Extending STRATA’s predictions to other cell types, STRATA also predicted a relative increased proportion of astrocytes and oligodendrocytes in Disc1-NAC treated animals, which is consistent with extensive literature findings demonstrating the neuroprotective effect of NAC by protecting again oxidative/proteotoxic stress (*27*–*30*).

**Fig. 4.**
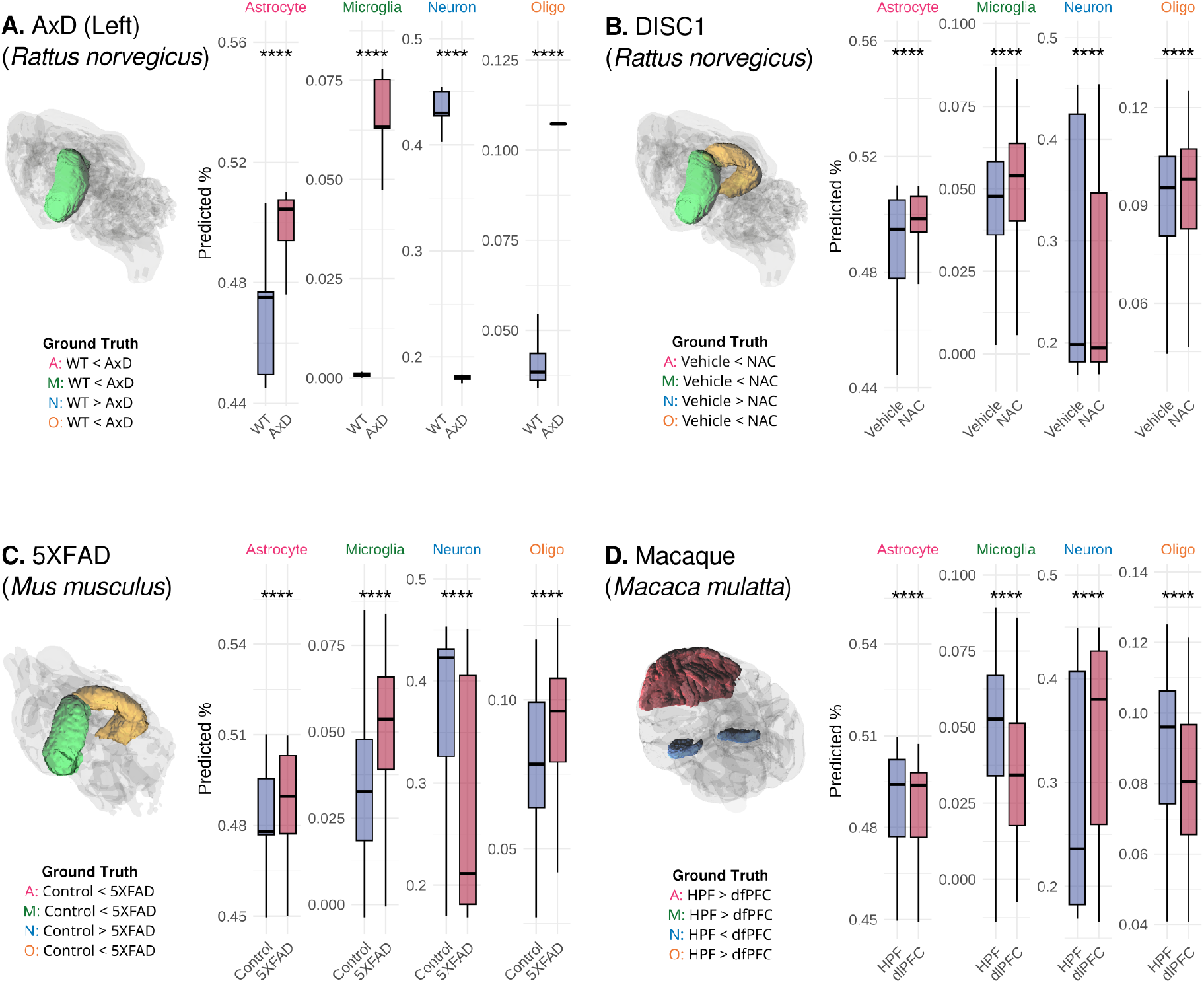
STRATA-predicted cell type proportions. Cell type populations were estimated across multiple test datasets representing various disease models, species, and ROIs. ROIs examined in each model are highlighted on accompanying 3D-glass brain volumes. Orange denotes the right hemisphere; green, left hemisphere except in the macaque dataset, where the hippocampal formation is shown in blue and the dorsal lateral prefrontal cortex is shown in red. Relative cellular populations were calculated in designated ROIs between disease/treatment groups and control animals. In the macaque experiment, relative differences in cell populations between the dorsolateral prefrontal cortex and the hippocampus were assessed. Test datasets: AxD (Left): Contralateral hippocampus of the Alexander’s disease (AxD) rat dataset used for training (training performed on right hippocampus only). Blue = wildtype; Red = AxD. DISC1: DISC1 knockout rat dataset, both hippocampi. All rats are DISC1 knockouts. Blue = vehicle treatment; Red = N-acetylcysteine (NAC) treatment. Macaque: Regional comparison in macaque dataset. Blue = hippocampal formations (HPF, left and right); Red = dorsolateral prefrontal cortex (dlPFC). 5XFAD: 5XFAD knockout mouse dataset, both hippocampi. Blue = control; red = 5XFAD knockout. Each subplot displays predictions for a single cell type, with the y-axis representing the predicted proportion. Box plots reflect the distribution of all predictions generated by the ensemble of XGBoost models for each cell type. Statistical comparisons were performed using the Wilcoxon rank-sum test and were Bonferroni corrected. Asterisks denote significance levels: \**p*<0.05, \*\**p*<0.01, \*\*\**p*<0.001, \*\*\*\**p* < 0.001; absence of asterisks indicates non-significance.

We next sought to determine the generalizability of the STRATA model. As STRATA is predicated on the relationship between brain cytoarchitecture and cell density, we reasoned that provided the known gross anatomical, cytoarchitectural, and functional conservation of the hippocampus across species (*31*), the STRATA model, which was trained on rat imaging data, should also perform well when analyzing NODDI imaging data of the hippocampus from other species. To test this hypothesis, using NODDI imaging data acquired from a 5XFAD transgenic mouse of Alzheimer’s disease (AD) (*32*), STRATA correctly predicted the relative increased proportions of astrocytes and microglia (*33*), the relative higher proportion of oligodendrocytes (manifesting as disease-associated oligodendrocytes) (*34*), and the relative decreased proportion of neurons present in the hippocampus of 5XFAD animals when compared to wild-type controls, the latter of which is a well-described neuropathologic correlated seen in the 5XFAD model (*35*) (**Fig. 4C**). To further test the predictive power of STRATA, we next used NODDI imaging data acquired from rhesus macaque monkeys (*M. mulatta*). Here, in the absence of an explicit well-characterized disease model, we used STRATA to predict relative differences in cell type proportions between the hippocampus and the dorsal lateral prefrontal cortex (dlPFC), which harbor significant differences cell type proportions than the hippocampus based on snRNA-seq data (*36*). Predicted cell type proportions of neurons, astrocytes, oligodendrocytes, and microglia from STRATA were compared against snRNA-seq transcriptomic data from these same regions of interest (*36*). snRNA-seq data from the dlPFC and the hippocampus demonstrate a relative higher percentage of neurons in the dlPFC compared to the hippocampus (dlPFC, 89.9%; hippocampus, 70.6%) and a relative decreased percentage of astrocytes, microglia, and oligodendrocytes in the dlPFC compared to the hippocampus (astrocytes [dlPFC, 4.4%; hippocampus, 8.5%], microglia [dlPFC, 0.27%; hippocampus, 3.98%], oligodendrocytes [dlPFC, 2.56%; hippocampus, 3.27%]), which align with the predicted relative cell type proportions from STRATA (**Fig. 4D**). Notably, an analysis of topological variable importance reveals that similar to how STRATA performed under training, STRATA exhibited strong feature independence and broad feature coverage with low feature covariance during prediction, which likely contributes to its strong predictive performance and the observed trans-species generalizability of the technique (**fig. S4**).

## Discussion

The STRATA model presented here is a new machine learning pipeline coupling spatially informed volumetric image quantification with quantitative neurobiology. This technique utilizes MRI data to enable the non-invasive estimation of relative differences in cell type proportions for neurons, astrocytes, oligodendrocytes, and microglia between groups. The ability of STRATA to deconvolve relative changes in cell populations from NODDI MRI builds on over a decade of progress in dMRI including advances in the mathematical and biophysical modeling of the dMRI signal, improved MR instrumentation, and recent efforts towards neurobiological quantification of the dMRI signal. The success of the STRATA model also stems from its ability to simultaneously address and overcome significant challenges in the clinical translation of qMRI. As qMRI techniques conventionally seek to accurately and reproducibly measure macromolecular properties of tissues, it is inherently susceptible to the instruments used to make these measurements and are thus confounded by system imperfections such as inhomogeneities in the static (*B*_0_) and applied radio frequency (*B*_1_) magnetic field and the lack of standardization in scanners and vendors leading to variability in measured quantitative parameters such as R1*, R2*, among others. Because STRATA quantifies the spatial relationship of imaging (intensity) data and not the measured quantitative MR values themselves (analogous to how clinical radiologists interpret imaging studies), STRATA is robust to the variability challenges present in qMRI. These inherent advantages are evident in our work with data acquired on different scanners with different field strengths, gradients, coils, and image acquisition protocols (4.7T Agilent vs 3.0 GE Magnus scanners) with no cross-vendor harmonization or scanner calibration performed prior to data collection or analysis. This out-of-the-box capacity to deconvolve relative differences in cellular proportions in rats, mice, and non-human primates in the hippocampus and in other gray matter structures like the dorsal lateral prefrontal cortex using only a small model built on rat hippocampal data further buttresses the robustness of the STRATA model and its trans-species generalizability. Interestingly, with STRATA demonstrating predictive power in other untrained and previously unseen gray matter structures like the dlPFC, this suggests STRATA is able quantify the relationship between gray matter topology and its intrinsic relationship to cellular populations akin to the converging representations seen in other deep learning models (*37*).

Our development of STRATA also intersects with the rising interest in quantifying the cellular landscape of the brain and introduces a novel non-invasive approach to quantify these changes to complement other single-cell and spatial -omic technologies. In the context of qMRI, STRATA addresses a longstanding challenge in qMRI by introducing a methodological approach to directly quantify the cellular neurobiology of the brain rather than through indirect imaging phenotypes that are both insensitive and non-specific. As a result, the ability to non-invasively resolve relative changes in cellular composition would have far reaching clinical implications including detecting acute inflammation (increased abundance of microglia), demyelination (oligodendrocyte depletion), and neurodegeneration (neuron loss) for pathologic diagnosis, treatment monitoring, and perhaps most importantly, inference into causal mechanisms of neurologic illness.

Understanding the neurobiological correlates of health and disease is a longstanding challenge in the basic and clinical neurosciences. STRATA enables direct non-invasive interrogation and quantification of the cellular dynamics of the brain to gain insight into the causal neurobiological mechanisms of disease. More broadly, the proof-of-principle application of TDA to NODDI dMRI that underlies the STRATA model is highly generalizable. Biological phenotyping of diagnostic imaging data has long been pursued to advance clinical care goals in precision diagnostics, therapeutics, and treatment monitoring. Notably, our machine learning pipeline can be flexibly applied to MRI, CT, PET, mammography, and other diagnostic imaging modalities to quantitatively phenotype diagnostic imaging data to enable specific imaging correlations to biological data to advance precision diagnostics in an era of personalized medicine.

## Acknowledgements

We thank past and current members of the Yu laboratory whose contributions, valuable discussions, and ideas provided the rationale and basis for the STRATA algorithm. We are grateful to David M. Holtzman, MD for providing insightful feedback and for his expertise and helpful discussions. We also acknowledge outstanding support of Michael Anderle and the Brain Imaging Core at the Waisman Center at the University of Wisconsin-Madison.

## Funding

Department of Radiology, University of Wisconsin School of Medicine and Public Health (JJY)

University of Wisconsin Science and Medicine Graduate Research Scholars (LAV)

National Institutes of Health grant T15LM007359 (MBF)

National Institutes of Health grant NS110719 (TLH)

National Institutes of Health grant F30MH140382 (RDR)

National Institutes of Health grant GM140935 (RDR)

Morse Society Fellowship (RDR)

## Author contributions

Conceptualization: LAV, JJY

Data curation: LAV

Funding acquisition: AMMS, DCD, TKU, JJY

Investigation: LAV, MBF, TLH, RDR, JMGG, APS, PCF, SAH

Methodology: LAV, MBF, JJY

Project administration: JJY

Resources: AMMS, DCD, JJY

Supervision: AMMS, DCD, JJY

Validation: LAV, JJY

Visualization: LAV, MBF, JJY

Writing-Original Draft: LAV, JJY

Writing-review & editing: LAV, MBF, TLH, RDR, JMGG, APS, PCF, SAH, AMMS, DCD, TKU, JJY

## Data and materials availability

All data and code to run the STRATA algorithm and to reproduce the major results of this manuscript are available in Dryad.

## Supplementary Materials

Materials and Methods

Supplementary Text

Figs. S1 to S4

Tables S1 to S3

References (38-51)

## Supplementary Materials

### Materials and Methods

#### MRI Acquisition and Preprocessing

*Ex vivo* imaging and analysis, including standard data preprocessing, study template generation, and region of interest (ROI) analyses, was performed as previously described *(18)*. In brief, all rats and mice were transcardially perfused with PBS followed by 4% paraformaldehyde (PFA) in PBS. The brains were then extracted from the cranial vault and post-fixed in 4% PFA. Brains were maintained at 4°C in 4% PFA in PBS until washing 2 × 24 hours with PBS followed by transfer into a custom-built holder immersed in Fluorinert (FC-3283; 3M, St. Paul, MN) and imaged with a 4.7T Agilent magnetic resonance imaging system with a 3.5-cm-diameter quadrature volume RF coil (Agilent Technologies, Santa Clara, CA). Multislice, diffusion-weighted spin echo mages were used to acquire 10 non-diffusion-weighted images (b = 0 s/mm2) and 75 diffusion-weighted images (25 noncollinear, diffusion-weighting directions: b=800 s/mm2; 50 noncollinear, diffusion-weighting directions: b=2000 s/mm2). Diffusion imaging was performed with echo time/repetition time = 24.17/2000 ms, field of view = 30 × 30 mm2, and matrix = 192 × 192 reconstructed to 256 × 256 over 2 signal averages. Raw data files were converted to NIFTI format, and FSL was used to correct for eddy current artifacts. FSL output volumes were converted to NIfTI tensor format for use with the DTI-TK software package. DTI-TK was employed to estimate a study-specific tensor template, to which subject tensor volumes were spatially normalized. Multi-shell diffusion data were fit with the Microstructure Diffusion Toolbox. An additional compartment of isotropic restriction was included to account for potential fixative effects as recommended . For rats, a DTI-based adult rat brain atlas (UNC-Chapel Hill) was used as a template to define ROIs, the left and right hippocampus and neocortex. For mice, a DTI-based adult mouse brain atlas was used as a template to mask out the left and right hippocampus *(38)*. Age, cohort size, and voxel resolution are summarized in Table S1.

**Table S1.** Summary of age, cohort size, and voxel resolution for rat and mouse datasets.

| Dataset | Species | Age | $n$ | Resolution | Citation |
| --- | --- | --- | --- | --- | --- |
| AxD | Rat<br>*Rattus norvegicus* | 56 days | Wildtype, Male = 6;<br>Wildtype, Female = 6;<br>AxD, Male = 6; AxD,<br>Female = 6 | Scanned at either<br>0.35 mm × 0.25 mm<br>× 0.25 mm or 0.25<br>mm isotropic; all<br>resampled to 0.35 mm<br>isotropic | [1] |
| DISC1 | Rat<br>*Rattus norvegicus* | P50 | All DISC1; Female =<br>6; Male = 6 | Voxel size of 0.25 mm<br>× 0.25 mm × 0.35<br>mm | [9] |
| 5XFAD | Mouse<br>*Mus musculus* | P55 | All Male; Control = 6;<br>5XFAD = 6 | 0.25 mm isotropic | Unpublished |

All procedures involving non-human primate tissue were conducted in compliance with protocols approved by the Institutional Animal Care and Use Committee of the University of Wisconsin and in accordance with NIH guidelines. At the time of necropsy, brains were perfused with 4% paraformaldehyde (PFA) and post-fixed for more than two weeks. For long-term preservation, tissues were subsequently transferred to 30% sucrose in phosphate-buffered saline (PBS). One day prior to imaging, samples were equilibrated in PBS at 15-21°C, and PBS removed immediately before imaging. No gross neuropathological abnormalities were detected in any specimens included in this study.

Two female Rhesus macaque (*Macaca mulatta*) brains were included in the study, aged 2.9 and 3.7 years postnatal, respectively. Magnetic resonance imaging (MRI) were obtained using a 3 T MAGNUS system (GE Healthcare, Waukesha, WI) and a 32-channel phased array receive coil (Nova Medical, Wilmington, MA). The MAGNUS platform delivers maximum gradient amplitudes of 300 mT/m and slew rates of 750 T/m/s using the standard clinical 3.0 T system power electronics (Signa Premier/Architect, GE Healthcare, Waukesha, USA). Data included multi-shell diffusion-weighted MRI (dMRI) acquired with a single-shot spin-echo EPI sequence at 1 mm isotropic resolution (slice thickness = 1 mm, acquisition matrix = 128 × 128), with in-plane acceleration factor = 2 and simultaneous multi-slice factor = 2. To help reduce geometric EPI distortions, the acquisition used a coronal prescription with the sample positioned in the sphinx position, resulting in the anterior portion of the brain facing the back of the bore (superior in scanner coordinates). Phase encoding was applied along the left–right axis. Imaging parameters included TE = 42 ms, TR = 4.52 s, flip angle = 90°, receiver bandwidth = 3906.25 Hz/pixel ( ≈ 256 *µ*s dwell time), effective echo spacing = 277.8 *µ*s, and total readout time = 35.28 ms.

For each nonzero b-value shell – 350 s/mm^2^ (24 volumes), 1000 s/mm^2^ (48 volumes), 2000 s/mm^2^ (72 volumes), 3000 s/mm^2^ (128 volumes), and 4000 s/mm^2^ (192 volumes) – an even number of volumes was acquired, with half collected using one phase-encoding direction and the other half using the reversed polarity, enabling robust susceptibility distortion correction while maintaining consistent diffusion encoding.

For the b = 0 s/mm^2^ shell (21 volumes total), the second phase-encoding series included one additional volume compared to the first, ensuring all diffusion-encoding directions across series were unique for a finer overall angular sampling. Additionally, b = 0 s/mm^2^ volumes were interleaved with diffusion-weighted volumes throughout each series to monitor and correct for temporal signal drift. The total acquisition time was approximately 18 min each of the series, yielding a combined total scan time of roughly 36 min for the complete reversed-PE dataset (485 volumes). See Tables S2 and S3 for full scanning parameters.

**Table S2.** Shell volume counts by b-value and phase-encoding direction for macaque data.

| b-value (s/mm<sup>2</sup>) | Volumes total | Volumes (PE dir A) | Volumes (PE dir B) |
| --- | --- | --- | --- |
| 0 | 21 | 10 | 11 |
| 350 | 24 | 12 | 12 |
| 990 | 48 | 24 | 24 |
| 2020 | 72 | 36 | 36 |
| 3010 | 128 | 64 | 64 |
| 4050 | 192 | 96 | 96 |

**Table S3.** Key imaging parameters for macaque data.

| Parameter | Value |
| --- | --- |
| Scanner | GE SIGNA MAGNUS 3T |
| Sequence | Single-shot spin-echo EPI |
| Internal sequence name | EPI2 |
| Voxel size | 1 mm isotropic |
| Slice thickness / spacing | 1 mm / 1 mm |
| Matrix | 128 × 128 |
| Acceleration (in-plane / SMS) | R=2 / MB=2 |
| Prescription | Coronal |
| Phase-encoding | Left-right |
| TR / TE | 4.52 s / 42 ms |
| Flip angle | 90° |
| Receiver bandwidth | 3906.25 Hz/pixel ($\sim 256 \mu\text{s}$ dwell time) |
| Effective echo spacing | 277.8 $\mu\text{s}$ |
| Total readout time | 35.28 ms |
| SAR | 0.38 W/kg |
| Total acquisition time (series A) | 18.23 min |
| Total acquisition time (series B) | 18.31 min |
| Total acquisition time (both) | 36.54 min |

The dMRI data were corrected for eddy currents, sample motion, susceptibility-induced geometric distortions, thermal noise, and Gibbs ringing artifacts. A brain mask was generated using a combination of ITK-SNAP’s manual and automated segmentation tools. This mask was used to remove signal resulting from the saline solution suspending the sample. Diffusion tensor imaging (DTI) and neurite orientation dispersion and density imaging (NODDI) parameter maps were subsequently computed, with all modeling steps accounting for gradient non-linearities in the prescribed diffusion encodings. To account for tissue fixation effects in the ex vivo preparation, the NODDI fixed parameters for parallel intracellular diffusivity and free water compartment diffusivity were adjusted to 0.2 *µm*^2^ × *ms*^−1^ and 2 *µm*^2^ × *ms*^−1^, respectively. These values were chosen based on rough estimates of the axial diffusivity in white matter and the mean diffusivity in ventricular spaces, respectively.

Following parameter map generation, image-header coordinates were corrected to match scanner coordinates using a combination of FSL’s fslorient -deleteorient, fslswapdim, and fslorient -setqformcode commands. The fractional anisotropy (FA) map was then diffeomorphically aligned to the ONPRC18 template using ANTs. The inverse of the resulting transformations was applied to the template labels to bring them into the native diffusion space, enabling extraction of scalar measures across the template-defined white and gray matter regions.

#### Topological Data Analysis

Topology is the mathematical study of objects through their connectivity and the spaces they enclose, which are robust to warping and deformation but not cutting and gluing *(39)*. Recent mathematical developments have led to the emergence of topological data analysis (TDA), the application of topological methods to analyze complex and large datasets *(40)*. One of these methods is homology, which associates topological features with a sequence of algebraic objects called homology groups, denoted *H*_*k*_. *H*_0_ contains all the connected components in an object, such as points, lines, or geometric faces in an object. *H*_1_ contains all the 1-holes in an object, empty two dimensional space enclosed by some *H*_0_ loop. *H*_2_ then contains all the 2-holes, empty three dimensional space enclosed by some *H*_0_ loop, and so on. One of the central techniques in TDA is persistent homology, which tracks the evolution of objects in these homological groups across different spatial and/or temporal scales *(41)*. More detail is available in the Supplementary Text.

First, each image is converted into a point cloud embedded in 4D space, with dimensions *X,Y*,*Z* – the spatial coordinates of voxels – and *I*, the intensity of a voxel. In order to not emphasize one dimension more than the other, we introduce min-max scaling for each dimension. We can then start to draw edges between points and build topological components called simplices. A simplex is a generalization of a triangle to both lower and higher dimensions. A 0-simplex is an individual point, a 1-simplex is a line, a 2-simplex is a solid triangle, and a 3-simplex is a solid tetrahedron. Each simplex is formed by the connection of lower-dimensional ones, starting from the 0-simplex, and each simplex is a single *H*_0_ object. A collection of simplices is a simplicial complex. In particular, the simplicial complex formed by connecting points in a dataset that are within some distance of each other is a Vietoris-Rips (VR) complex.

A VR complex is the set of all simplexes, *S*, formed by connecting a finite set of points 𝒫 given all their composing vertices are within some chosen distance 2*ϵ* from each other,

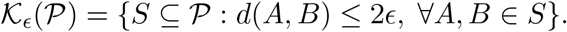

The distance metric *d* is 4-dimensional Euclidean distance,

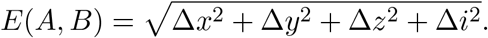

This is computed by forming closed-ball neighborhoods of radius *ϵ* around each point in the point cloud,

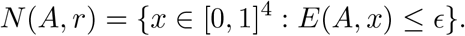

If there are two neighborhoods that overlap, *N* (*A, r*) ∩ *N* (*B, r*) ≠ ∅, the edge *E*(*A, B*) = 2*ϵ* is drawn. Using these neighborhoods, we can also determine whether an enclosed space exists and whether or not it is filled.

Instead of any optimal choice of *ϵ*, we consider a sequence of thresholds starting at *ϵ*_0_ = 0, then increased by some chosen step size *e* such that *ϵ*_*i*_ = *ϵ*_*i*−1_ + *e* for index *i >* 0, and ending at *ϵ*_*t*_ = *t*. Here, we choose *t* = max {*E*(*A, B*)}. By creating the respective VR complex, 𝒦_*i*_, for each *ϵ*_*i*_, we can build a nested sequence of simplicial complexes,

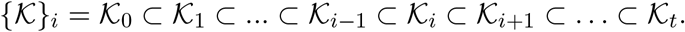

A hole may then first appear at some 𝒦_*b*_ and persist until it is closed at some 𝒦_*d*_ where *ϵ*_*b*_ *< ϵ*_*d*_. From the perspective of the closed-ball neighborhoods, at *K*_*i*_ we first draw an *H*_0_ loop that encloses some space and at *K*_*j*_ the neighborhoods completely overlap and cover it. By tracking the birth and death of objects in each of our homological groups across our sequence of thresholds, called a filtration, we can quantify the topology of a dataset from multiple scales, in a local-to-global manner.

Each feature that is born and dies can be matched with an interval [*ϵ*_*b*_, *ϵ*_*d*_) that tracks its birth and death times. A stack of these intervals is called the persistence barcode *(42)*. A common way to represent these intervals is the persistence diagram, which graphs the birth time of some feature on the x-axis and the death time on the y-axis. However, since *ϵ*_*b*_ *< ϵ*_*d*_ always, all points in this graph appear above the diagonal. Many vectorization methods have been developed in order to convert a persistence barcode or diagram into a vector that is usable in machine learning. One common approach is the persistence image *(43)*. First, a persistence diagram is linearly transformed to instead plot [*ϵ*_*b*_, *ϵ*_*d*_ − *ϵ*_*b*_), removing the appearance of a diagonal. The yaxis now then represents lifetime or persistence, rather than death. Next, a normalized symmetric Gaussian distribution is created around each point [*ϵ*_*b*_, *ϵ*_*d*_ − *ϵ*_*b*_) in the transformed diagram. A persistence surface function, the sum of all the normalized Gaussian distribution is then defined, in order to create an integrable surface over the same range as the transformed persistence diagram. By integrating across *b* equally spaced bins on each axis, we obtain a *b* × *b* matrix, with each entry being a new feature that represents the overall activity in a respective part of the diagram. One thing to note is that *H*_0_ components are all born at 0 but die at unique times. Therefore, the vectorization result of this dimension is instead a vector of size *b*. More information is available in the Supplementary Text.

This was done through giotto-ph, a multi-core implementation of the Ripser algorithm for VR persistent homology in Python *(44)*. Each image was processed through the Center for High Throughput Computing at the University of Wisconsin-Madison. For computational feasibility, a sparse approximation approach was used to simplify the input point cloud. This uses a greedy permutation in order to find the most representative edges for the topology of the point cloud, and introduces a hyperparameter that indicates the level of simplification *(45)*. For all images, this was chosen to be 1. Conversion of persistence diagrams, the output of giotto-ph, to persistence images was done through the giotta-tda package in Python *(46)*. This introduces two more hyperparameters, one for the number of bins and another for the standard deviation of the normalized symmetric Gaussian distributions. For all images, the standard deviation was chosen to be 0.001 and the number of bins was chosen to be 8. This provides 136 features for each image, 64 features from each 8 × 8 persistence image for *H*_1_ and *H*_2_ and another 8 from the 8 × 1 persistence image for *H*_0_.

Given there are two MRI modalities, NDI and ODI, for each subject imaged there are 272 features used for training and prediction.

#### Bulk RNAseq Preprocessing

Bulk transcriptomics samples were obtained from previously published data *(17)*, and were collected as follows. Males and females of each genotype were sacrificed at 8 weeks of age by CO2 asphyxiation (N = 4 per group, 16 samples total), and the right hippocampus was collected immediately on ice and frozen (∼ 80°C) before subsequent processing. The hippocampus included Ammon’s horn, the dentate gyrus (DG), and the subiculum. The alveus was used as a dorsal boundary to separate the hippocampus from the subcortical white matter, and parahippocampal regions were removed from the ventral/lateral boundaries. Excess white matter from the fimbria and fornix were also removed. RNA was extracted with TRIzol reagent per the manufacturer’s protocol (Invitrogen, Thermo Fisher Scientific) and treated with TURBO DNase (TURBO DNA-free Kit, Ambion, Thermo Fisher Scientific), and RNA integrity was determined with an Agilent 4200 TapeStation system for quality control. Libraries were prepared from 1 *µ*g total RNA with integrity numbers (RIN) between 8.4 and 9.3 for sequencing with TruSeq Stranded mRNA Sample Preparation kit (Illumina). Libraries were quantified with PicoGreen reagent (Thermo Fisher Scientific) and assayed with an Agilent TapeStation system to confirm integrity before sequencing with an Illumina NovaSeq X Plus (2 × 150bp, 70 M reads per sample, two lanes). Base calling was performed using Bcl2fastq (v2.20.0.422), read trimming with Skewer *(47)*, read alignment with STAR *(48)*, and expression estimation with RSEM *(49)*.

#### Cellular Deconvolution

In order to estimate the proportion of celltypes found in a bulk RNA seq dataset, the DWLS method was used *(20)*. The implementation of DWLS was done in R using the omnibus package omnideconv *(50)*. In this method, an annotated single cell RNAseq dataset is needed to generate representative gene profiles of each cell type expected to be found in the tissue. In order to best match the bulk data, we used a publicly avaliable dataset of single cell transcriptomics of the dorsal dentate gyrus. The rats sampled were 60 day-old, compared to 8 weeks old in our bulk data, and contained both control animals and those treated with fluoxetine *(51)*. After deconvolution, each bulk sample has an estimated percentage of each cell type in the original single cell reference. These included microglia, inhibitory neurons, excitatory neurons, astrocyte subtype 1, oligodendrocyte subtype 1, oligodendrocyte subtype 2, mossy cells, endothelial cells, oligoprogenitor cells (OPCs), granule cells, and berg glia. Due to the small percentage of some of these cell types, and in order to make this as generalizable to other regions as possible, we simplified our analysis. Subtypes were condensed together – such as for microgila, neurons, and oligodendrocytes – while certain celltypes were labeled as Other and excluded (mossy cells, endothelial cells, berg glia, OPCs, and granule cells). As there was only one subtype for astrocyte avaliable in the dorsal dentate gyrus scRNAseq dataset, this was renamed to astrocyte. See Figure S1 for more detail.

**Fig. S1.**
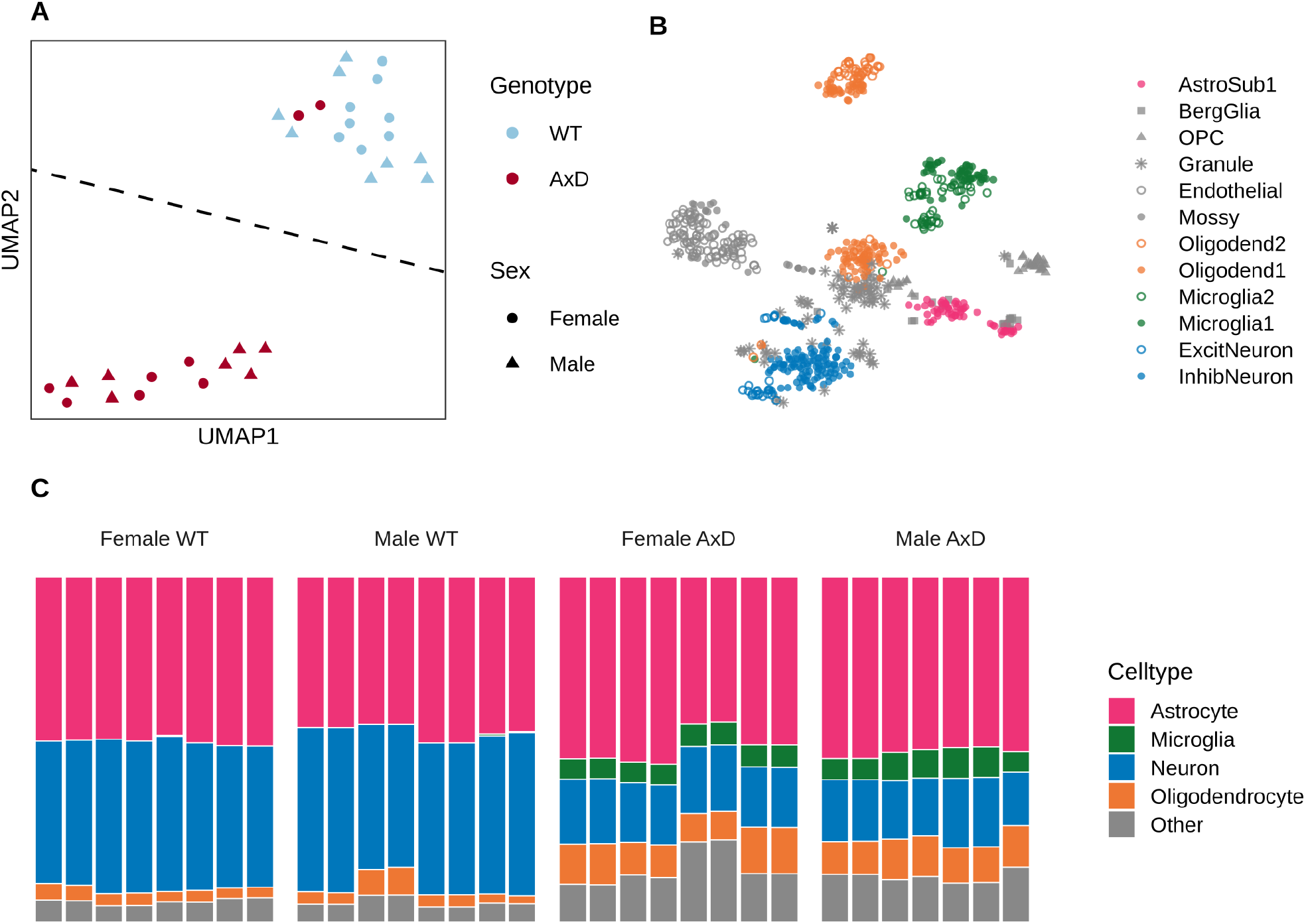
**(A)** Bulk RNA-seq data (UMAP projection shown) was deconvolved using a reference single cell RNA-seq dataset (tSNE projection shown) via the Dampened Weighted Least Squares (DWLS) method, yielding estimated cell type proportions for each sample. Bulk RNA-seq data is from the right hippocampus of wildtype or Alexander Disease (AxD) genetic model rats (male and female) **(B)** scRNAseq reference data is from the dorsal dentate gyrus of similarly aged rats. **(C)** After deconvolution, subtypes of major cell types (astrocyte, microglia, neuron, and oligodendrocytes) were grouped together and cell types with small percentages, or that are hippocampus-specific, were grouped together as Other. Bar plots show the deconvolved percentages of these cell types for each rat in the bulk RNA-seq dataset.

#### XGBoost

Training data comes from two cohorts: one that was used for *ex vivo* imaging and another that was used for bulk transcriptomics, which was deconvolved into representative cell types as described above. Each cohort contains animals that are either male or female and either wildtype (WT) or the Alexander’s disease genetic model (AxD). In the transcriptomics cohort, bulk RNA sequencing was only available from the right hippocampus. Therefore, in the imaging cohort only topological features of the right hippocampus were considered for training. Features were min max scaled within each dataset tested, and within each ROI for the macaque and mice datasets. Additionally, the macaque scans were upsampled by a factor of three to yield point clouds with a comparable number of points to those obtained from the rat and mouse datasets. In order to pair data from different cohorts a permutation approach was used. Each rat from the imaging cohort was paired with all rats in the transcriptomics cohort of both the same sex and the same genotype, in order to create multiple paired samples. This acknowledges the fact that since we can’t define an optimal pairing for each rat, we must consider all biologically-plausible pairings. In addition, this bootstraps our dataset and provides more samples to train on.

In R, XGBoost regression models were trained seperately for each cell type, including astrocytes, neurons, microglia, and oligodendrocytes, as to test the efficacy of predicting each cell type in isolation *(21, 22)*. Each model was trained for 250 boosting iterations using a squared-error objective function and root mean squared error (RMSE) as the evaluation metric. Additionally, we adopted a leave-one-out (LOO) approach where all pairs including a specific subject were removed during training in order to not rely on any specific sample from either cohort, and a separate XGBoost model was made for each LOO trial. Thus, many XGBoost models were made for each cell type and we rely on ensemble predictions from all LOO models. As ensemble predictions were not normally distributed, the Wilcoxon Rank Sum test was used to compare whether samples came from different distributions and were Bonferroni corrected. Variable importance features, including frequency, gain, and cover from XGBoost, as well as SHapley Additive exPlanation (SHAP) values were used to interpret model results *(24)*. See Figure S4 for more detail.

**Fig. S2.**
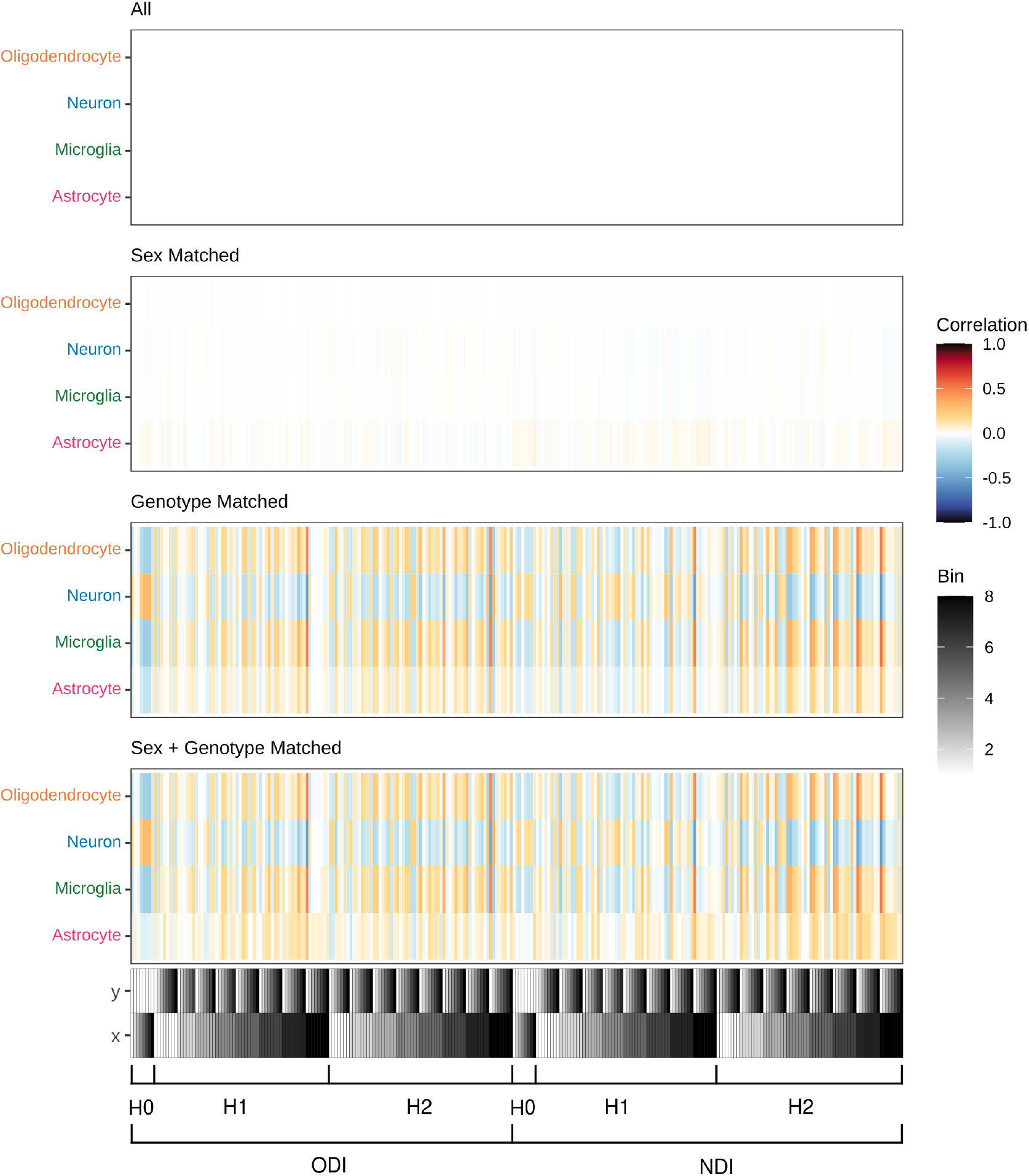
Permutation Heatmaps. Heatmaps show pairwise Pearson correlations between deconvolved cell types from the bulk RNA-seq training cohort, y-axis, and topological data analysis (TDA) features from paired rats in the imaging cohort, x-axis. Tile color indicates direction and magnitude of correlation (red = positive, blue = negative). Each subplot corresponds to a different pairing schema used to match RNA and imaging data: All; All possible pairings considered. Sex-matched; Only rats of the same sex were paired, regardless of genotype (wildtype or AxD). Genotype-matched; Only rats of the same genotype were paired, regardless of sex. Sex + Genotype matched; Only biologically coherent pairings, matching both sex and genotype, were included. The bars below all heatmaps indicate the origin of the imaging feature being correlated. X and Y denote bin positions within the 8×8 persistence image, represented using a grayscale (1 = white, 8 = black). Additional labels specify the homological dimension (H0, H1, H2) and the imaging modality (NDI, ODI).

**Fig. S3.**
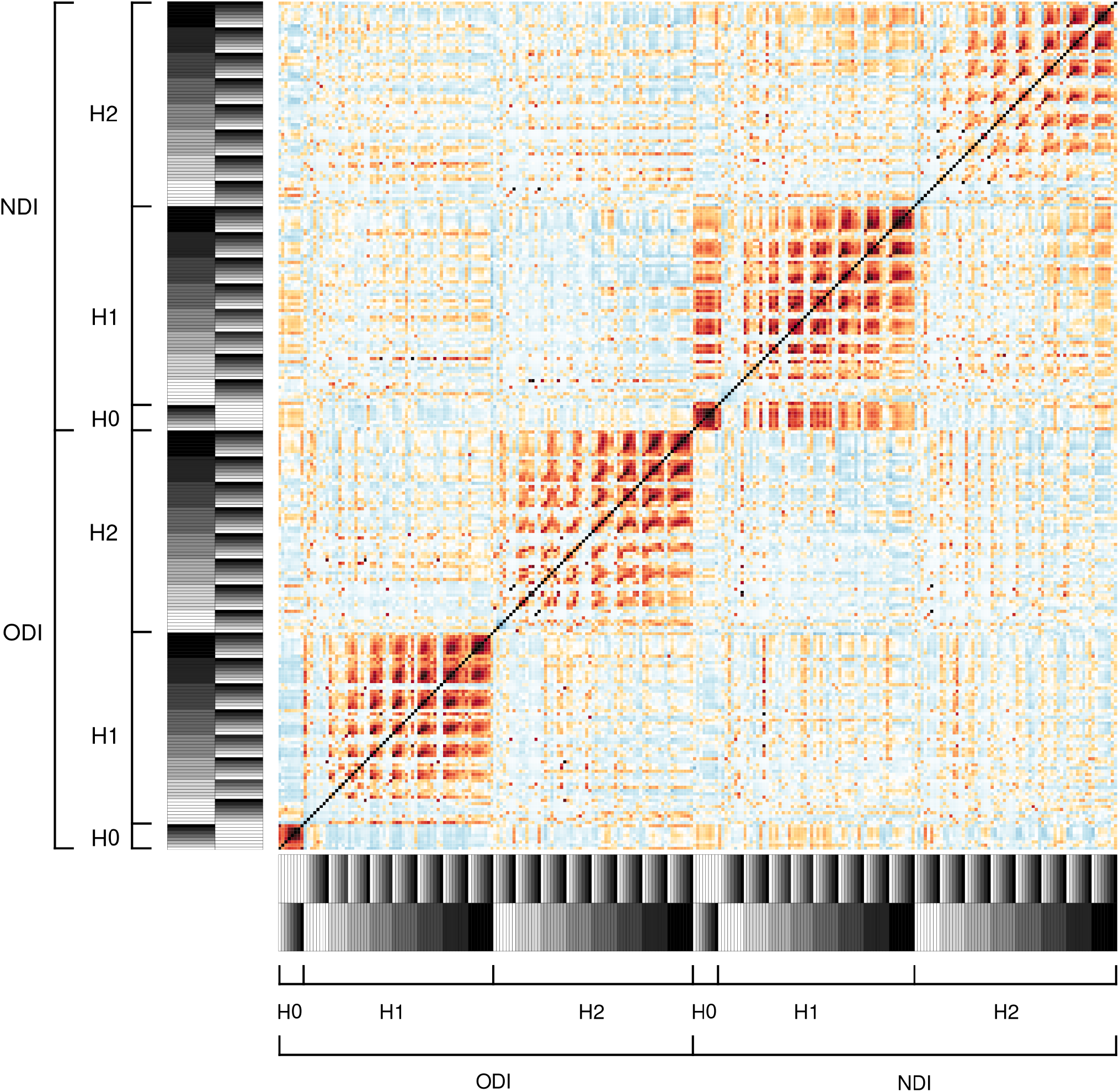
Correlation Matrix for TDA Features in Training Data. Symmetric matrix showing pairwise Pearson correlations between different persistence image features obtained from topological data analysis (TDA). Tile color indicates direction and magnitude of correlation (red = positive, blue = negative). Bars along both the x and y axes indicate the origin of the imaging feature being correlated. X (leftmost along y-axis, bottommost along x-axis) and Y (rightmost along y-axis, upmost along x-axis) denote bin positions within the 8×8 persistence image, represented using a grayscale (1 = white, 8 = black). Additional labels specify the homological dimension (H0, H1, H2) and the imaging modality (NDI, ODI).

**Fig. S4.**
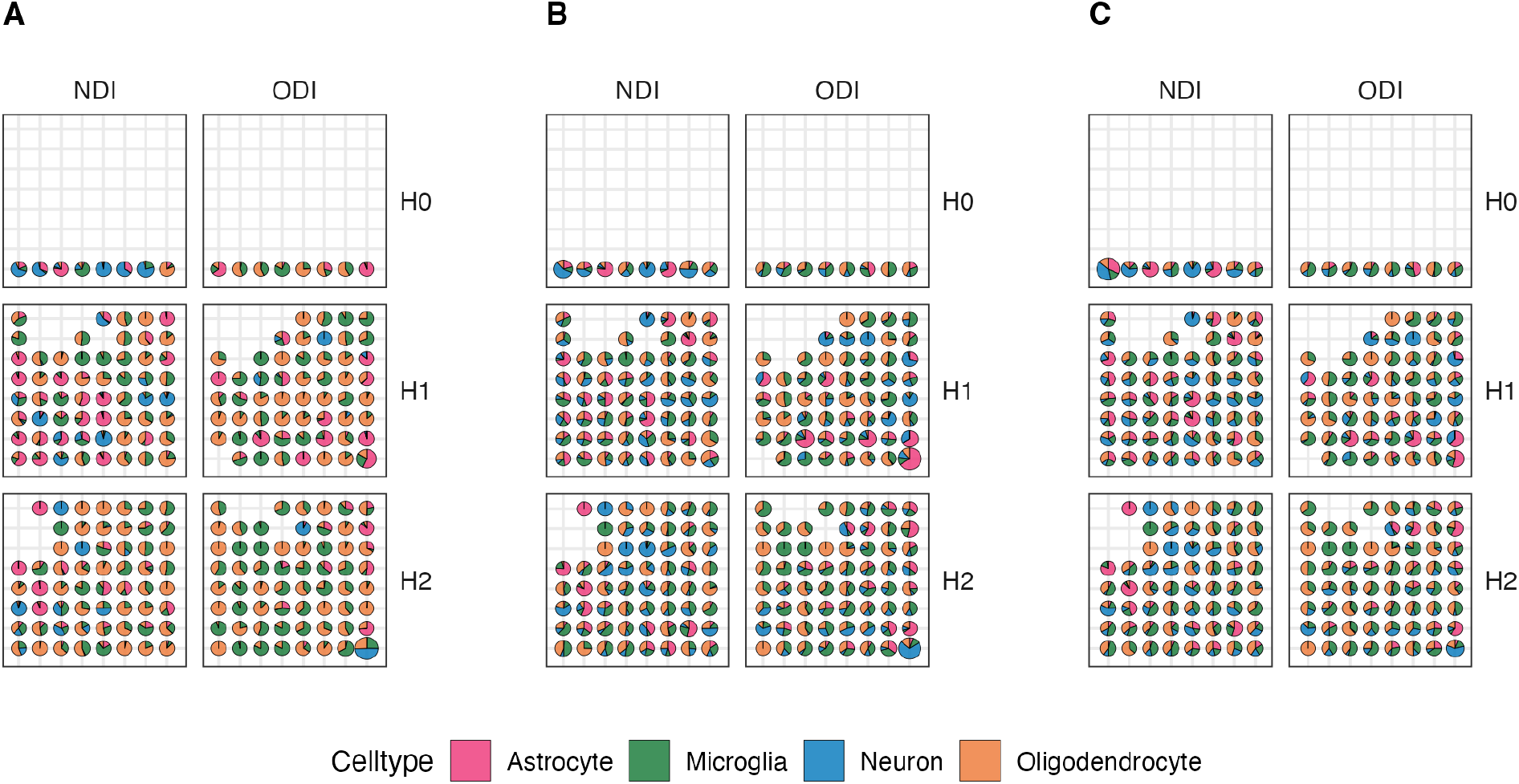
Grid of plots showing the mean values for features derived from persistence image from different variable importance metrics. Each plot corresponds to the persistence image of a specific modality (columns) and homological dimension (rows). Within each plot, the x-axis represents birth time and the y-axis persistence, corresponding to bin positions in the 8×8 persistence image. Each pie chart represents a single persistence image feature and shows its mean contribution to the prediction of different cell types, calculated across the ensemble of XGBoost models. The size of each pie chart reflects the overall importance of that feature, with larger charts indicating higher mean values. Bins without a pie chart represent features that were not used in the final models. **(A)** Mean Gain: improvement in model accuracy brought by decision tree splits involving the feature, indicating how helpful a feature is in reducing prediction error. **(B)** Mean Cover: number of samples affected by decision tree splits involving the feature, indicating how broadly the feature is used across the dataset. **(C)** Mean Frequency: number of times a feature is used to split data across all trees in the model, indicating the feature is frequently considered useful during tree construction.

Because subjects in the imaging cohort do not have ground truth cell-type proportions and the only available ground truth comes from the transcriptomics cohort, individual-level prediction accuracy was not evaluated. Matching predicted values across hemispheres in the imaging cohort is invalid, as the right hippocampus was only used to generate training pairs and lacks ground truth itself. Similarly, comparing predictions for the left hippocampus to ground truth from the right is ill-posed. Therefore, standard metrics like MAE or R2 are not applicable in this cross-cohort design. Instead, our evaluation focuses on group-level inference and whether predicted distributions align with known biological trends that can be verified in the dataset with immunohistochemistry or other methods.

### Supplementary Text

Point Cloud Construction Each image is first converted into a point cloud embedded in 4D space, 𝒫. Some voxel *A* in the image can now be considered a node in 𝒫 with the domain *D* ⊂ *R*^4^ where *A* = {*x*_*A*_ ∈ *X, y*_*A*_ ∈ *Y, z*_*A*_ ∈ *Z, i*_*A*_ ∈ *I*} and *D* = {*X, Y, Z, I*} . *X, Y*, and *Z* are the spatial coordinates of the voxel itself, and *I* corresponds to it’s intensity, commonly visualized as the color of the voxel. In order to not emphasize one dimension more than the other, we introduce min max scaling, 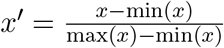, for each dimension in our domain. From this we obtain the new scaled point cloud 𝒫^′^ with the domain *D*^′^ = [0, 1]^4^ ⊂ *R*^4^ . *A* ∈ 𝒫 now has a corresponding *A*^′^ ∈ 𝒫^′^ where 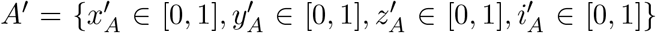. This is done for each image individually. Between the voxel *A*^′^ ∈ 𝒫^′^ and another *B*^′^ ∈ 𝒫^′^ we can draw an edge *E*(*A*^′^, *B*^′^) such that *E* is the Euclidean distance between these two points,

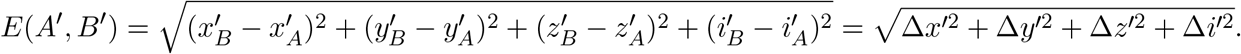

This can be interpreted as a measure of dissimilarity, considering both macro-structure (through *X, Y, Z*) as well as micro-structure (through *I*). For simplicity, in the rest of the section we will assume all 𝒫 are already min max scaled.

In the unscaled point cloud, each spatial dimension has a unit step between adjacent voxels of 1. However, after scaling this unit step is changed for each dimension independently such that they may no longer be equal. Additionally, it is impossible for any two points to share the exact same spatial position so *E*(*A, B*) *>* 0. There is no unit step for *I*, but there is a min(Δ*i*) and a max(Δ*i*) such that 0 ≤ min(Δ*i*) ≤ max(Δ*i*) *<* 1. Δ*i* = 0 if and only if the voxels share the exact same intensity. Each MRI image of a given species is registered to an atlas such that they have the exact same number and location of voxels, prior to conversion into point clouds. Here, the only difference between two point clouds will be in *I*. However, as the 4D embedding considers *X, Y, Z* and *I* equivalently, the underlying Euclidean distances between points will still be different. When comparing point clouds from different regions or species altogether, each of the spatial dimensions may lie in entirely different domains as well. By introducing min max scaling, we can ensure that all point clouds share the same domain *D* = [0, 1]^4^.

#### Topology and Homology

Topology studies objects based on their connectivity and the multidimensional spaces these components enclose, which are robust to warping but not cutting or gluing. This is done through the classification of objects by their Betti numbers, *β*_*k*_. *β*_0_ counts the number of connected components in an object, such as lines, disconnected points, and geometric faces. *β*_1_ counts the number of 1-holes in an object, or regions of 2D space enclosed by a *β*_1_ loop. *β*_2_ then counts the number of 2-holes in an object – regions of 3D space enclosed by a *β*_1_ shell. Two objects are homological to each other if they have the same *β*_*k*_ numbers, even if these objects have different geometries. A simple example of this is comparing a hollow circle with a hollow triangle. Although both of these objects are geometrically distinct, they both have *β*_0_ = 1 and *β*_1_ = 1 and are equivalent in topology. In order to calculate the Betti numbers of some embedded point cloud, we can build edges between points and start to build topological components called simplices.

A *k*-simplex is a *k*-dimensional polytope (a geometric object with flat sides) with *k* + 1 vertices. The 0-simplex is a single disconnected point, with a single vertex. The 1-simplex is a line, with two vertices as end points, formed by connecting at least two 0-simplices. The 2-simplex is a solid triangle, with three vertices and a 2D face, and can only be formed by connecting at least three 1-simplices. The 3-simplex is the solid tetrahedron, with four vertices and one 3D face, formed by the connection of at least four 2-simplices. The Betti numbers of all the simplices are *β*_1_ = 1, as they are all fully connected and contain no empty space. *β*_*>*1_ *>* 0 if and only if an enclosed space exists, such as in the hollow triangle or the hollow tetrahedron. A hollow triangle would have *β*_0_ = 1 and *β*_1_ = 1 as previously illustrated, while a hollow tetrahedron would have *β*_0_ = 1, *β*_1_ = 0, and *β*_2_ = 1 – as the *β*_0_ shell encloses a 3D volume and not a distinct 2D surface. A set of multiple simplices is a simplicial complex, denoted by 𝒦, under the condition that every face of some simplex in 𝒦 is itself a unique simple and any shared face between simplices is a component of each of them. The topology of a point cloud 𝒫 can then be calculated by connecting points to build simplices and calculating the *β*_*k*_ numbers of the constructed simplicial complex.

A VR complex is the set of all simplexes *S* formed by connecting a finite set of points *X* – defined in a metric space *M* – given all their composing vertices are within some chosen distance 2*ϵ* from each other,

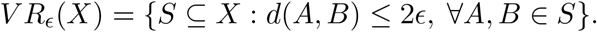

For a min-max scaled 4D point cloud, *M* = [0, 1]^4^, *X* = 𝒫, and the distance metric *d* is 4-dimensional Euclidean distance,

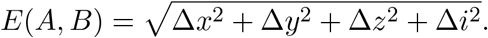

Therefore, a triangle, line segment, or tetrahedron is included in the complex if and only if all the edges between it’s vertices are ≤ 2*ϵ*. This is determined by forming closed-ball neighborhoods of radius *ϵ* around each point in the point cloud,

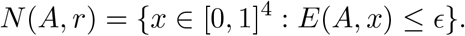

If there are two neighborhoods that overlap, *N* (*A, r*) ∩ *N* (*B, r*) ≠ ∅, the edge *E*(*A, B*) = 2*ϵ* is drawn and a 1-simplex is built.

A 1-hole in a VR complex is then built by at least three points *A, B*, and *C* such that their pairwise neighborhoods overlap to build a *β*_0_ loop that encloses a 2D space. That is,

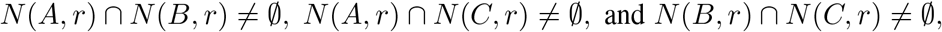

but,

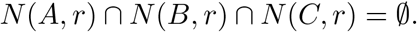

If a sufficiently large *ϵ* is chosen then *N* (*A, r*) ∩ *N* (*B, r*) ∩ *N* (*C, r*) ≠ ∅, and this is now a 2-simplex; the *β*_1_ hole no longer exists and the face is filled in. By comparing the *ϵ* when a hole is first formed or ‘born’, *ϵ*_*b*_, and the time of it’s closure or death, *ϵ*_*d*_, the size of the enclosed space can be estimated. This logic extends as well as to higher *β*_*>*1_.

By progressively increasing *ϵ* and constructing a new VR complex each time, we can track the birth and death of *β*_*k*_ components and obtain a more complete view of the topology. At *ϵ*_0_ = 0 the simplex is a point cloud with every point disconnected, a collection of *p* 0-simplices where *p* is the number of points in 𝒫. This may then be called 𝒦_0_ = *V R*_0_(𝒫), where *β*_0_(*K*_0_) = *p* and *β*_*>*1_(*K*_0_) = 0. *ϵ* is then increased by some chosen step size *e* such that *ϵ*_*i*_ = *ϵ*_*i*−1_ + *e* for index *i >* 0. At *ϵ*_*i*_, we then obtain _*i*_ = *V R*_*i*_(𝒫).

As points are connected throughout this process, *β*_0_ continuously decreases as the original 0-simplices connect together to give rise to 1-simplices, holes, and higher-order simplices. Since all *β*_0_ components are born at *ϵ*_0_, the elder rule states that the vertex appearing first in a list of points is the one that persists. For example, if 0-simplices *A* and *B* where to merge to form the 1-simplex *AB*, it would be the component representing *A* that would continue, while the one representing *B* would die. The corresponding *β*_0_ numbers here would drop from 2 to 1. Importantly, the elder rule only applies to *β*_0_, because *β*_1_ and *β*_2_ holes do not merge throughout the filtration and instead are individually opened and closed. Some maximum distance *ϵ*_*t*_ must be chosen to end this process, with a common choice being the maximum *E* distance possible between any two points in 𝒫. This creates a nested sequence,

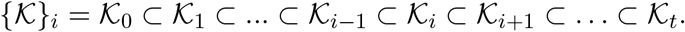

This entire process is called the Vietoris-Rips filtration, and the resulting nested sequence of simplicial complexes is called a filtered simplicial complex. Tracking the lifetime, or persistence, of topological components in a dataset in relation to some filtration process is the definition of the persistent homology technique in topological data analysis. In order to computationally do this, we turn to algebra.

#### Algebra

A *k*-chain in a simplicial complex 𝒦 is the linear combination of a finite number of *k*-simplices, *S*_*k*_ ∈ 𝒦, that are not necessarily connected,

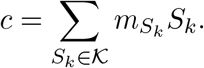

The algebraic field used for computations is *F*_2_ = {0, 1} and all calculations are in modulo 2. The coefficients *m*_*i*_ then are always either {0, 1} and are used to encode whether a simplex is included in a chain, *m* = 1, or excluded, *m* = 0. If a simplex is included twice in a chain, modulo 2 ensures that it is canceled out. *C*_*k*_(𝒦) is the set of all *k*-chains in 𝒦. From this we can define the boundary map,

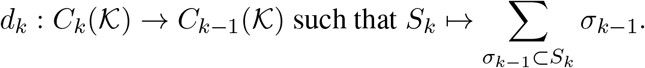

This map deconstructs some *k*-simplex *S*_*k*_ into the sum of it’s lower dimensional components, *σ*_*k*−1_, which must be (*k* − 1)-simplices themselves. The composition of this map *d*_*k*−1_ ° *d*_*k*_ always maps to 0 as the boundaries of simplices themselves cannot have boundaries.

The kernel set,

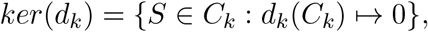

consists of all the elements of *C*_*k*_ that map to the identity element 0, the simplices in 𝒦 that do not have a boundary. The image set,

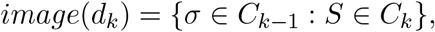

contains all the boundary elements created from the map. It can be shown that *image*(*d*_*k*+1_) ⊂ *ker*(*d*_*k*_). The usefulness of this can best be shown through an example. For a 2-simplex *ABC*, it follows that 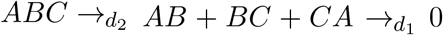, so *AB, BC, CA* ∈ *image*(*d*_2_) and *AB, BC, CA* ∈ *ker*(*d*_1_). Assume instead we had a chain of 1-simplices that formed a hollow triangle *c* = *AB* + *BC* + *CA*. We can then say 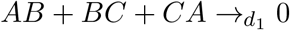 so *c* ∈ *ker*(*d*_1_). However, *c* ∉ *image*(*d*_2_) as it is not the boundary of a 2-simplex and cannot even be sent through *d*_2_. Therefore *c* ∈ *ker*(*d*_*k*_)*/image*(*d*_*k*+1_), the set of all elements that are in the kernel but not the image.

We therefore define the quotient space

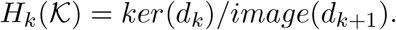

This space holds all the topological components in the *k* dimension, which are all homologous to one another, and thus is called a homological group. *H*_0_ contains all the connected components in 𝒦, *H*_1_ contains all 1-holes in 𝒦, and *H*_2_ contains all the 2-holes in 𝒦. The formal definition of a Betti number is actually *β*_*k*_(𝒦) = *rank*(*H*_*k*_(𝒦)), the number of objects in the corresponding homological group. Let *n* denote the dimension of 𝒦, meaning that 𝒦 holds simplices of dimension 0 to *n*, but no simplices of dimension *k > n*. It then follows that *H*_*k>n*_(𝒦) = 0 and *C*_*k>n*_(𝒦) = 0. We can now build a sequence,

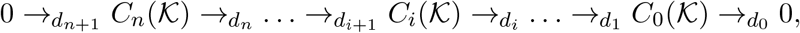

which we will call the chain complex. From this chain complex, we can then calculate the *H*_*k*_(𝒦) for all *k* ∈ {1, …, *n*}.

In order to track components in each of our *H*_*k*_ groups across the filtered simplicial complex 𝒦_0_ ⊂ … ⊂ 𝒦_*i*_ ⊂ … ⊂ 𝒦_*t*_, and record when they appear and disappear, we need to create a mapping between different complexes *f* : 𝒦 → 𝒦^′^ from which we can build a corresponding mapping *h*_*k*_ : *H*_*k*_(𝒦) → *H*_*k*_ (𝒦^′^). This is possible because simplicial homology is a functor, a mapping between categories of mathematical objects that is structurally-preserving. We can define *f* simply as the inclusion map, which maps the same elements to themselves in the next step of the filtration. Any *f* : 𝒦 → 𝒦^′^ map we make then induces another inclusion map,

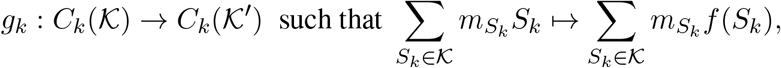

where *f* (*S*_*k*_) ∈ 𝒦^′^. In other words, given a map *f* between these simplicial complexes there exists a map *g*_*k*_ that maps any *k*-chain in 𝒦 to its corresponding *k*-chain in 𝒦^′^ through *f* . Moreover, this map respects the boundary mapping such that 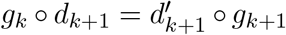, where *d*_*k*+1_ is the boundary map for *K* and 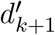 is the boundary map for *K*^′^. This ensures that boundaries only map to boundaries and the chain complex structure is preserved. The mapping *g*_*k*_ itself then induces another inclusion map *h*_*k*_ : *H*_*k*_(𝒦) → *H*_*k*_ ( 𝒦^′^) where *c* ↦ *g*_*k*_(*c*).

This chain of maps *f* → *g* → *h* is fundamental for capturing the birth and death of holes throughout 𝒦_0_ ⊂ … ⊂ 𝒦_*i*_ ⊂ … ⊂ 𝒦_*t*_. Suppose we have a 1-hole in 𝒦_*i*_ formed by the 1-simplices *AB, BC*, and *CA*, and in 𝒦_*i*+1_ the hole is closed such that *ABC* is now a 2-simplex. Through *f* we have *ABC* ∈ 𝒦_*i*_ ↦ *ABC* ∈ 𝒦_*i*+1_ and through *g* we have *AB* + *BC* + *CA* ∈ *C*_1_( 𝒦_*i*_) ↦ *AB* + *BC* + *CA* ∈ *C*_1_( 𝒦_*i*+1_). Notice that *AB* + *BC* + *CA* ∈ *image*(*d*_2_ (𝒦_*i*+1_)) as it is the boundary of the 2-simplex in 𝒦_*i*+1_, so it follows *AB* + *BC* + *CA* ∉ *H*_1_( 𝒦_*i*+1_). However, *AB* + *BC* + *CA* ∈ *H*_1_( 𝒦_*i*_). Therefore, when we induce *h*_1_ : *H*_1_( 𝒦_*i*_) → *H*_1_( 𝒦_*i*+1_) we must have *h*_1_(*AB* + *BC* + *CA*) ↦ 0 by necessity. This is the definition of death in persistence homology, when we can no longer map the hole to itself and must instead map to 0.

In addition, functoriality says that that the induced functions are stable for chain of mappings 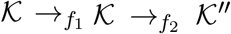. We can then say that for a filtered simplicial complex the inclusion map *f*_*i,j*_ : 𝒦_*i*_ → 𝒦_*j*_ induces 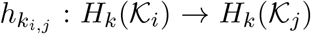 for any *i, j* ∈ {0, 1, …, *t*} such that *i* ≤ *j*. It is now valid to construct a chain of inclusion maps 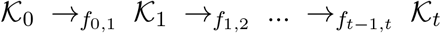 that induces another chain of inclusion maps 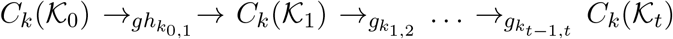, which in turn induces 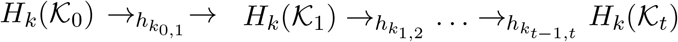. The set of homological groups and their mappings to each other,

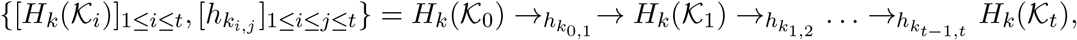

is formally called the *k*-th persistence module, which carries the information of birth and death for all the holes in *H*_*k*_(𝒦) throughout the filtration.

Let *𝓁* = *β*_*k*_{𝒦}_*i*_, the total number of *H*_*k*_ components that ever existed throughout our filtration. For any persistence module there exists a unique (up to reordering) set of intervals 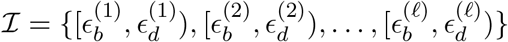 where 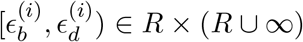 such that the persistence module decomposes into

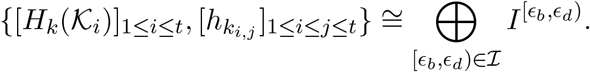

In other words, each of those holes and its maps is matched to a collection of vector spaces called the interval module,

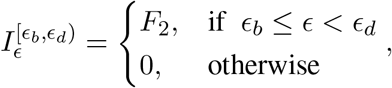

alongside their own internal inclusion mappings 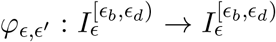 for *ϵ*_*b*_ ≤ *ϵ* ≤ *ϵ*^′^ *< ϵ*_*d*_ . This lets us continuously and consistently track the lifespan of this hole as an interval [*ϵ*_*b*_, *ϵ*_*d*_) ∈ ℐ, where *ϵ*_*b*_ is the radii that formed 𝒦_*b*_ – the simplicial complex where the hole first appeared in our filtration – and *ϵ*_*d*_ the radii that formed 𝒦_*d*_ – the simplicial complex when that hole dies. The direct sum, ⨁, is simply a stack of all these interval modules, defined on the same range as our filtration, [0, *ϵ*_*t*_], such that each *I* lives independently from each other. The reason we define 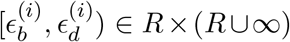 is to handle cases when a hole still survives at the end of our filtration, such that the interval for that hole would be [*ϵ*_*b*_, ∞ ). This set of intervals ℐ is the bar-code representation of the holes in our point cloud 𝒫 – a complete summary of the local to global topology – and can now be statistically interpreted or vectorized for machine learning tasks.

#### Vectorization

A common way to represent 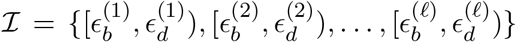 is as a persistence diagram, where each tuple is graphed onto a 2D plane with an x axis for birth time and the y axis for death time. However, as *ϵ*_*b*_ *< ϵ*_*d*_ always, none of these tuples can appear below the diagonal. By instead plotting birth against lifetime, *ϵ*_*d*_ − *ϵ*_*b*_, this effect can be removed, which is more convenient for vectorization. Formally, this transformation is defined as *T* : *R*^2^ → *R*^2^ such that *T* (*x, y*) = (*x, y* − *x*). This gives a new set of intervals 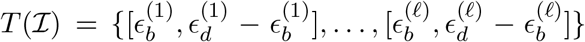. However, *H*_0_ here is unique. We always start our filtration with *p H*_0_ components, which are all born at 0 but die at unique times. Therefore, *𝓁* = *p* and 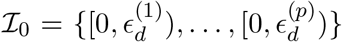, so a 1D representation is sufficient. Furthermore, ℐ_0_ = *T* (ℐ_0_) as 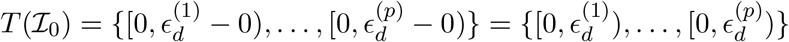.

In general, the cardinality of ℐ_0_ may be different than ℐ_1_ and ℐ_2_ for some filtered simplicial complex, {𝒦*}*_*i*_. In addition, the cardinality of ℐ_*k*_ for some 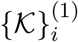 may differ from another 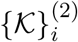. Therefore, in order to compare ℐ_*k*_ for different 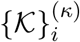, one must create a robust and equivalent vectorization such as a persistence image. First, we construct a normalized symmetric Gaussian distribution for each *u* = (*x*_*u*_, *y*_*u*_) ∈ *T* (ℐ), such that

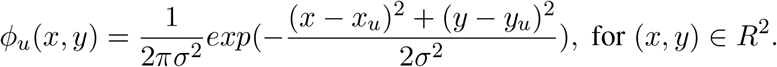

The persistence surface is then the function,

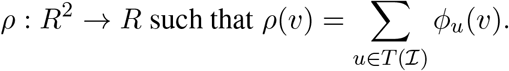

This gives a density kernel based on the sum of overlapping Gaussian distributions created from each point in *T* (ℐ). We can then then take the integral across both axes,

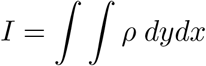

in order to calculate the kernel density for a region of x and y values. For the x axis we then define an equal partition 𝒳 = {𝒳_0_, …, 𝒳_*b*_*}* where 𝒳_0_ = 0 and 𝒳_*b*_ = max{*x*_*u*_ : (*x*_*u*_, *y*_*u*_) ∈ *T* (ℐ)*}*, and for y similarly create 𝒴 = {𝒴_0_, …, 𝒴_*b*_*}*. These partitions then create a set of *b* − 1 bins for each axis, such as {[𝒳_0_, 𝒳_1_], …, [𝒳_*b*−1_, 𝒳_*b*_]*}* for the x axis. Therefore, the density at some *x* × *y* bin is then,

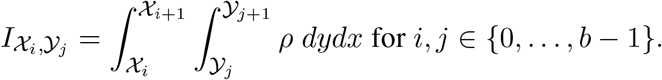

For *H*_0_, this is a vector of size (*b* − 1),

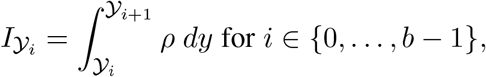

and the Gaussian kernel is replaced with it’s 1-dimensional counterpart. This creates a matrix of size (*b* − 1) × (*b* − 1) for each ℐ_*k>*0_,

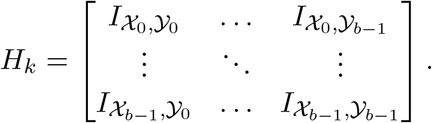

For ℐ_0_ this is a vector of size (*b* − 1),

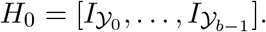

These matrices then provide a way to statistically compare the topology from a point clouds which vary both in their original number of points *p* as well as their Betti numbers in each dimension.

